# Scaling the Dynamics of Coiled Coils

**DOI:** 10.64898/2026.07.21.739852

**Authors:** Prithwidip Saha, Yogesh Saravanan, Arin Marchesi, Claire Valotteau, Patrick Revy, Tanya T Paull, Salvatore Assenza, Karl-Peter Hopfner, Mauro Modesti, Felix Rico

**Author notes:** Present address: Institute of Physics, University of Augsburg, Universitätsstrasse 1, 86159 Augsburg, Germany (Prithwidip Saha).

## Abstract

Coiled coils are structural motifs in proteins that play diverse functions. In MRE11-RAD50 (MR) complexes, ATP-driven changes in coiled coils are essential for DNA break sensing. However, coiled coil dynamics and its modulation by protein conformational changes remain unclear, partly due to the lack of quantitative tools. Here, we used high-speed atomic force microscopy (HS-AFM) for real-time visualization of the coiled coil conformational dynamics of individual MR complexes from bacteria and human homologs, and a biomedically relevant variant. The mean square deviation of the end-to-end distance of the coiled coils revealed a power-law scaling with time, conserved across conformational states, homologs, and variants, suggesting a universal dynamic scaling. Coiled coils behave as semi-flexible filaments with strong internal friction, leading to relaxation times that were seconds-long and varied among conformational states and variants. Molecular dynamics simulations indicated that strong friction arose from long-lifetime contacts between coils. Our results suggest that MR complexes modulate the coiled coil dynamics to mediate long-range allosteric and *allodynamic* communication during DNA repair.

## Introduction

Coiled coils are conserved and versatile structural motifs of proteins (1, 2). Often consisting of heptad repeat sequences, they are formed by two or more interwinding α-helices. Coiled coils play multiple structural and functional roles in biological systems, serving as molecular spacers, scaffolding large protein complexes, mediating vesicle tethering, and acting as mechanosensors to transmit conformational changes for motor activity (3–5). Moreover, they have also emerged as orthogonal building blocks for the precise control of protein interactions and cellular function (6). Variation in the sequence and length of coiled coils across protein systems is expected to lead to differences in the flexibility and, importantly, the dynamic response, thereby influencing their functional roles.

During DNA recognition and cleavage, highly dynamic coiled coil domains are essential structural components of a family of protein complexes known as structural maintenance of chromosomes (SMC). In SMC complexes, the conformational dynamics of cohesin and condensin have been characterized, providing key insights into their DNA-loading mechanism (4, 5, 7, 8). An evolutionarily conserved DNA repair factor related to SMC complexes is the human MRN complex (hMRN), which consists of three proteins: MRE11, RAD50, and NBS1. RAD50 homologs feature a coiled coil region being 15- to 60-nm-long, depending on the species (9, 10), which is essential to support genome integrity during MRN nucleolytic processing of DNA ends and subsequent DNA repair (11–15).

In bacteria, the respective homologs of MRE11 and RAD50, SbcD and SbcC, form the SbcCD complex, which processes palindromic DNA hairpins and cruciform DNA structures and endonucleolytically cleaves DNA strands near protein obstructions (16–19). The protein complex exists as a heterotetramer, comprising two SbcCs and two SbcDs. SbcC features an intramolecular antiparallel coiled coil region, with an ATP-nucleotide binding domain (NBD) forming, together with an SbcD dimer, the catalytic head domain at one end and a cysteine-based chelating motif forming the hook at the other end (20). Both the hook and the head regions facilitate SbcC dimerization (**Fig. 1A**). The catalytic head domain binds and cleaves DNA in a reaction involving conformational changes between the two NBDs induced by ATP. In these homologs (hMRN and SbcCD), the coiled coils act as a gate for DNA ends and as a mechanical device to regulate activation of the MRE11/SbcD nuclease within the catalytic head domain (3, 21–24). Further, coiled coils mediate long-range allosteric communication between the hook and the head domains (25) to regulate the DNA end engagement upon adopting different open and closed configurations. Therefore, the flexibility and dynamics of the coiled coil are expected to be important for its role as a structural gate for DNA ends.

**Figure 1:**
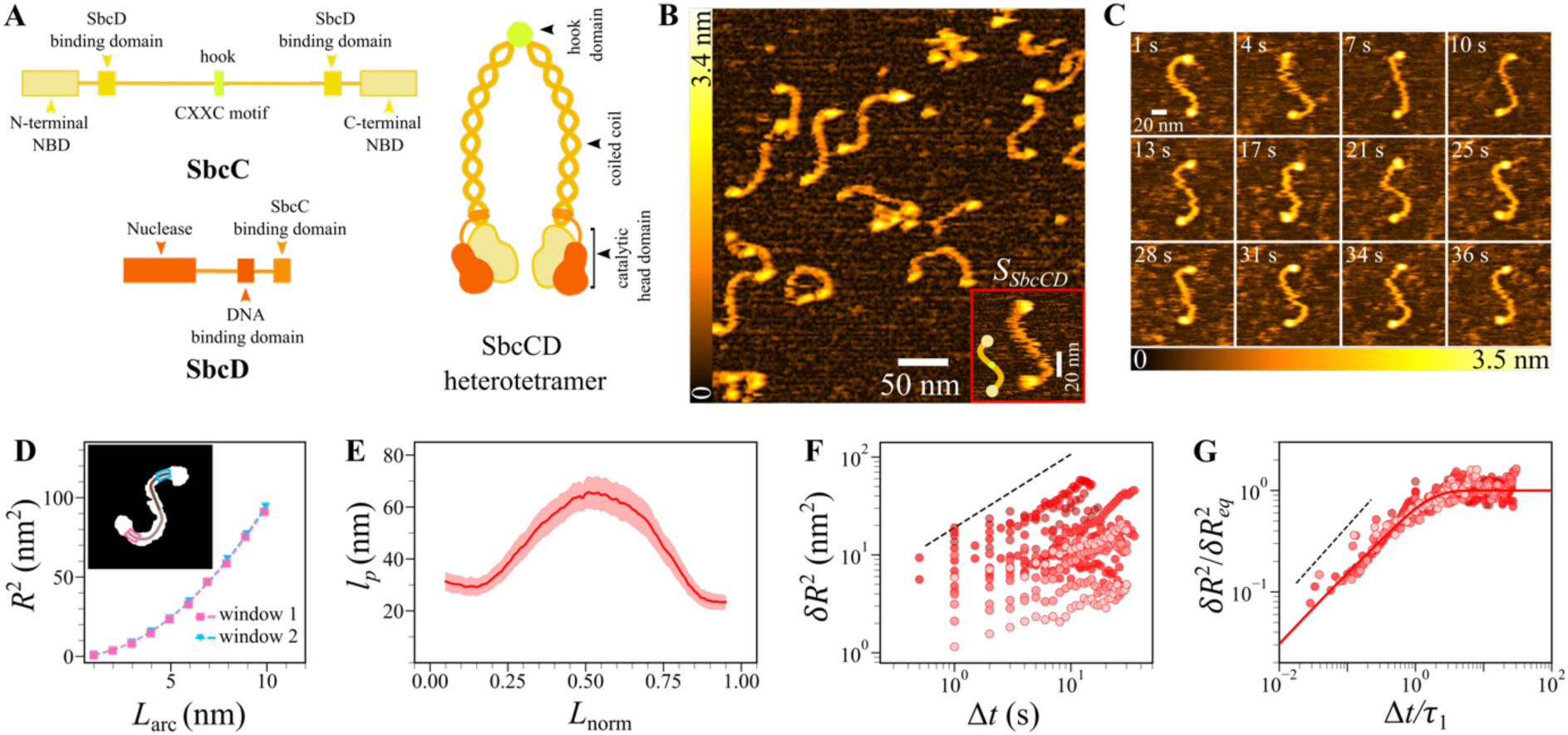
**A.** Left: Scheme of the domain organization of SbcC (yellow) and SbcD (orange). Right: Scheme of the SbcCD heterotetramer, a SbcC dimeric coiled coil linked by the hook and with SbcD bound to either head. **B.** HS-AFM topograph showing different conformations adopted by the SbcCD complex upon deposition on a mica surface. Inset: the most abundant *S* conformation with a schematic representation. **C.** Selected frames showing conformational dynamics of the dimeric coiled coil in *S*_*SbcCD*_. The timestamps are shown at the top left of each image. **D.** Squared end-to-end distance (*R*^2^) as a function of coiled coil arc length (*L*_*arc*_) measured at 1 nm steps within a ∼10 nm window (window 1: magenta with square, window 2: cyan with triangle) along each monomeric coiled coil segment (light and dark brown). The 2D WLC model (dashed lines) was fitted to the data to determine the persistence length *l*_*p*_. Inset: Binary image of the *S* conformation showing the interpolated skeletal coordinates of the dimeric coiled coil marked with light brown (monomeric segment 1) and dark brown (monomeric segment 2), excluding the globular head domains. **E.** Local persistence length *l*_*p*_ as a function of normalized contour length *L*_*norm*_ with shaded area representing SEM (*N*= 8 videos). **F.** Mean-square displacement of the end-to-end distance (*δR*^2^) for monomeric coiled coil segments for *S*_*SbcCD*_ as a function of elapsed time Δ*t* between HS-AFM frames. Shades of red in the data points represent different datasets from 8 HS-AFM videos, each point being a time-averaged value over all possible initial times. **G.** Plot of the rescaled MSD (*δR*^2^/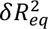) as a function of the rescaled time (Δ*t*/*τ*_1_), where 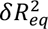 is the characteristic equilibrium saturation value and *τ* is the characteristic relaxation time. The solid red line is the predicted 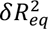 (**Eq. 1**). Dotted black lines in **F** and **G** are power laws of exponent 3/4.

The DNA binding and repair mechanism is highly dynamic and involves important conformational changes within the MR complex. Cryo-EM studies revealed high-resolution conformational snapshots of the head domain of SbcCD and hMRN complexes, revealing an ATP-bound resting conformation and a DNA-end recognition and cutting state (20, 26). Because of its high flexibility and dynamic nature, conformational changes have been inferred only from the proximal and distal coiled coil fractions (27, 28). Indeed, cryo-EM typically proposes mechanisms based on final observations, which may suffer from selection biases and obscure conformational flexibility. Conversely, atomic force microscopy (AFM) imaging in solution revealed high flexibility of the coiled coil regions in the MR complexes (29).

The coiled coil’s flexibility depends on both its sequence and the possible disruptions in the network of interactions between α-helices. Single-molecule force spectroscopy has been used to probe the mechanical response of coiled coils, including their unfolding and uncoiling under load (30–38). AFM snapshots of human MRE11-RAD50 further revealed heterogeneous flexibility along the coiled coil length (39). High-speed AFM (HS-AFM) has emerged as a powerful tool for visualizing the flexibility and dynamics of proteins, capturing real-time nanoscale conformational changes with sub-second time resolution (40, 41). Importantly, it allows quantifying the dynamics of proteins, for example, by determining the relaxation times of disordered regions (42). HS-AFM has reported various conformational states of coiled coil protein complexes, such as the golgins, laminins, archaeal MR and bacterial SbcCD complexes, revealing their highly dynamic structural flexibility (43–46). However, a quantitative understanding and a theoretical description of the structural dynamics of coiled coils are still missing (39, 41). Moreover, while essential to understanding allosteric function, how conformational changes in the complex affect the structural flexibility and dynamics of the coiled coils remains unexplored.

Here, we used HS-AFM imaging to visualize the structural dynamics of individual bacterial and human MR complexes in real time. HS-AFM videos displayed different conformational states and varying local flexibility among MR complex variants. Analysis of the end-to-end distance of the coiled coil revealed a power law response over time for all MR complex variants and conformations, suggesting universal scaling. Semi-flexible filament modelling allowed us to assign a fundamental relaxation time to each conformation and variant. The resulting seconds-long relaxation times suggested high internal friction. All-atom MD simulations confirmed the experimental scaling law and explained that the internal friction arises from the unusually long lifetime contacts between sliding coils. Our results suggest that the dynamic response of coiled coils is an overlooked parameter in understanding allosteric communication.

## Results

We obtained HS-AFM videos of the bacterial SbcCD complex weakly immobilized on a mica surface in physiological buffer. Topographs revealed complexes with different conformations (**Fig. 1B)**, where the coiled coils showed a height of ∼1.5 nm, inherent curvature, dynamic flexibility, and mobility via thermal fluctuations, while globular head domains located at both termini were higher (∼3 nm). The most abundant (∼70 %) conformation featured an S-shape (**Fig. 1B, C)**. We named this conformation as *S*_*SbcCD*_ based on its structural shape adopted by the deposited protein complex, and this naming convention is applied hereafter to all other conformations. The rarely found *Z*-shaped conformation (i.e., *Z*_*SbcCD*_, a mirror image of *S*_*SbcCD*_) indicated a preferred interaction of the head domains with the surface (Supplementary Figure **S1**). All observed conformations allowed the coiled coils to move freely on the surface. The movement of coiled coils along the fast and slow scanning axes of imaging showed similar behaviour, suggesting minor perturbation by the HS-AFM probe (Supplementary Figure **S2**). The average contour length of the coiled coil region was ∼90 nm, in agreement with the expected dimensions of two SbcC coiled coils connected via their hook region. While the mica-deposited complexes were expected to present both SbcC and SbcD, the volume of the head domains indicated that most complexes likely lack SbcD, therefore existing only as dimers of SbcC (Supplementary Figure **S3**). While rare, we also observed monomeric and multimeric forms of the protein complex, the latter bound by both the hook and head domains (Supplementary Figure **S4**). Some HS-AFM topographs revealed high-contrast features along the coiled coils, including one in the middle corresponding to the hook and two others near each head domain. AlphaFold2 predictions of the coiled coil segment suggest that these protrusions are due to the presence of kinks or local deformations in the coiled-coil assembly due to discontinuities of the heptad repeats (Supplementary Figure **S5**)(47).

To quantify the local flexibility of the coiled coils, we determined the persistence length (*l*_*p*_) along the contour length *L* (see Methods). For that, we extracted the coiled-coil skeleton of each molecule (per video frame), excluding the terminal globular domains (**Fig. 1D inset**) (48). Using a 10-nm moving window (rolling every 1 nm), we fitted the 2D worm-like chain (WLC) model to the measured squared end-to-end distance ⟨*R*^2^⟩ versus the window arc length *L*_*arc*_(**Fig. 1D)** (49). The fitted *l*_*p*_ values were then averaged over hundreds of frames and different molecules along the normalized contour length *L*_*norm*_. This analysis revealed that the local flexibility varies along the coiled coil length (**Fig. 1E**), ranging from ∼30 nm to ∼70 nm, with increased rigidity (higher *l*_*p*_) in the central region (hook). Previous AFM data on human MR complexes agreed with our results (39). The sharp change from low to high *l*_*p*_ (and high to low) coincided with the protrusion near the head observed in high-resolution topographs and a recurring high curvature region predicted by AlphaFold2 (Supplementary Figure **S5**) (47).

Next, we quantified the dynamics of each coiled coil segment of protein complexes from different HS-AFM videos. For that, we determined the mean square deviation (MSD) of the end-to-end distance ⟨*δR*^2^⟩ over the elapsed times (Δ*t*), from ∼0.5 to ∼100 s (see Methods). The resulting ⟨*δR*^2^⟩ as a function of Δ*t* revealed two main asymptotic regimes: a power law at short times and a plateau at longer times (**Fig. 1F**). Interestingly, the power law revealed subdiffusive motion (exponent <1), differing from the dynamics of an ideal chain (exponent =1). This dynamic response resembled that observed in semiflexible filaments, modelled by solving the Langevin equation for an elastic rod of contour length *L*, effective persistence length 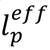, and friction coefficient per unit length *ζ*

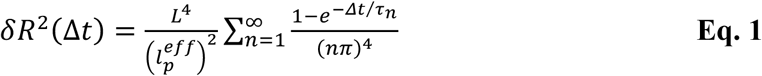

where *τ*_*n*_ is the characteristic relaxation time of the *n*-th bending mode of the filament (50) (see Methods). For each video and coiled coil, we fixed *L* to the average contour length and fitted **Eq. 1** to the MSD to extract 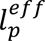 and *ζ* (Supplementary Figure **S6**). We then used these fitted parameters to rescale the dynamic data, normalizing the MSD to the characteristic equilibrium saturation value 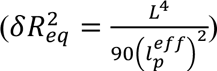 and the elapsed time to the characteristic equilibration time 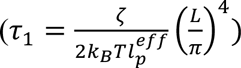, i.e., the relaxation time of the fundamental, slowest mode (*n*=1) (**Fig. 1G**, see **Table 1**). The rescaled data collapsed into a master curve, a power law at short times with an exponent of 3/4 (Supplementary Text **S3**) saturating to 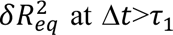. As expected from geometrical considerations, the MSD of the angle between the two ends of each coiled coil followed a power law of exponent 1/4 with the elapsed time (Supplementary Figure **S7**), although without clearly reaching equilibrium. This behaviour has been observed before in the context of actin filaments, although of tens-of-µm contour length (51, 52).

**Table 1.** Contour length *L*, effective persistence length 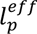, friction coefficient ζ, and relaxation time *τ* across conformations for coiled coils of SbcCD, SbcC, hMRN, hMR, hMR^E1035Δ^, hR and hR^E1035Δ^ derived from data analysis of MSD over time (mean ± SEM, with N=2*–*8 being the number of videos, each consisting of 24–70 frames). The conformational abundance was determined from overall AFM topographs presenting multiple complexes.

| Deposited protein complex | Conformation | Abundance (%) | $L$ (nm) | $l_p^{eff}$ (nm) | $\zeta$ ($10^{-3}$ pN·s/nm<sup>2</sup>) | $\tau$ (s) |
| --- | --- | --- | --- | --- | --- | --- |
| **SbcCD** | $S_{SbcCD}$ | 70 | $44 \pm 0$ | $59 \pm 6$ | $66 \pm 19$ | $7.2 \pm 2.5$ |
| | $C_{SbcCD}$ | 8 | $49 \pm 1$ | $68 \pm 6$ | $22 \pm 4$ | $2.6 \pm 0.7$ |
| | $braided_{SbcCD}$ | 4 | $55 \pm 2$ | $77 \pm 17$ | $15 \pm 6$ | $2.4 \pm 0.9$ |
| | $closed_{SbcCD}$ | 6 | $55 \pm 3$ | $65 \pm 3$ | $6 \pm 2$ | $1.1 \pm 0.6$ |
| | $closed_{SbcCD}^{ATP}$ | 9 | $54 \pm 2$ | $92 \pm 8$ | $14 \pm 4$ | $1.5 \pm 0.3$ |
| | $S_{SbcCD}^*$ | 8 | $42 \pm 2$ | $65 \pm 4$ | $32 \pm 8$ | $1.5 \pm 0.2$ |
| | $monomer_{SbcCD}$ | 1 | $42 \pm 3$ | $37 \pm 5$ | $6 \pm 1$ | $0.8 \pm 0.3$ |
| **SbcC** | $S_{SbcC}$ | 75 | $44 \pm 0$ | $75 \pm 7$ | $24 \pm 3$ | $1.6 \pm 0.2$ |
| **hMRN** | $open_{hMRN}$ | 13 | $57 \pm 2$ | $49 \pm 17$ | $34 \pm 16$ | $15.3 \pm 8.1$ |
| | $closed_{hMRN}$ | 87 | $51 \pm 1$ | $43 \pm 6$ | $22 \pm 7$ | $5.7 \pm 2.2$ |
| **hMR** | $open_{hMR}$ | 26 | $58 \pm 1$ | $59 \pm 14$ | $21 \pm 7$ | $10.3 \pm 4.4$ |
| | $closed_{hMR}$ | 74 | $57 \pm 1$ | $55 \pm 7$ | $11 \pm 2$ | $3.9 \pm 1.6$ |
| **hMR<sup>E1035A</sup> (hMRm)** | $open_{hMRm}$ | 52 | $48 \pm 1$ | $36 \pm 4$ | $15 \pm 3$ | $3.1 \pm 1.1$ |
| | $closed_{hMRm}$ | 48 | $59 \pm 2$ | $37 \pm 5$ | $6 \pm 1$ | $2.9 \pm 1.1$ |
| **hR** | $open_{hR}$ | 100 | $52 \pm 0$ | $40 \pm 6$ | $18 \pm 7$ | $4.7 \pm 1.8$ |
| **hR<sup>E1035A</sup> (hRm)** | $open_{hRm}$ | 44 | $54 \pm 3$ | $35 \pm 7$ | $29 \pm 14$ | $11.6 \pm 6.8$ |
| | $closed_{hRm}$ | 62 | $56 \pm 2$ | $32 \pm 3$ | $53 \pm 14$ | $18.9 \pm 4.5$ |

The effective persistence length of each monomeric segment (59±6 nm) obtained from the dynamic analysis fell within the range of the local persistence length from the local WLC analysis (30–70 nm). The average friction coefficient from all analysed videos was *ζ*_0_∼0.07±0.02 pN·s/nm^2^, leading to a characteristic equilibration time of a few seconds (*τ*_1_∼7±2 s). For complexes with a head domain volume corresponding to the expected SbcC+SbcD, the dynamic response was similar (Supplementary Figure **S3**), although saturating at shorter relaxation times (∼1.5 s, see 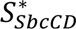 in **Table 1**). This suggests that binding of SbcD may induce a conformational change that affects coiled coil dynamics. Our results show that coiled coil segments in SbcCD complexes behave as semiflexible filaments, revealing subdiffusive motion. Importantly, the observed friction coefficient was exceptionally high. Indeed, for a free filament in water, one expects *ζ*_0_ = 4*πη*/ ln(2*L*/*r*) (50), where *η* ≃ 10^−9^ pN·s/nm^2^ is water viscosity and *r* ≃ 1 nm is the radius of the filament. For *L* ≃ 44 nm, *ζ*_0_ ≃ 2.5 · 10^−9^ pN·s/nm^2^ (corresponding to *τ*_1_ ≃ 0.2 μs), which is about seven orders of magnitude lower than our experimental value. Lubrication effects account only partially for this discrepancy, as according to the small-separation formula 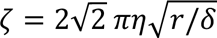 (53), one would obtain an increase of only an order of magnitude for *δ* ≃ 0.1 − 0.5 nm, i.e., substrate separation comparable to the size of a water molecule (54, 55). Similarly, the restricted water mobility close to the mica surface is expected to yield a larger effective local viscosity, yet typical values of hydrophilic surfaces correspond to at most an increase of an order of magnitude (56). The large increase is thus likely to emerge from intrinsic ruggedness of the bending free-energy landscape, leading to kinetic traps in the dynamics. Based on the classical derivation by Zwanzig for a one-dimensional potential (57), we estimate *ζ* = *ζ*_0_ exp[(*ϵ*/*k*_B_*T*)^2^], where *ϵ* is the energy scale of the landscape roughness; a value of *ϵ* ≃ 3.4 *k*_B_*T* would thus explain the residual discrepancy of five orders of magnitude. Microscopically, we identify two possible co-occurring mechanisms behind such ruggedness: heterogeneity of protein-mica interactions and internal friction of the coiled coil. In the former case, we speculate that the main source of heterogeneity is the non-uniform charge distribution along the coiled coil and mica (see Supplementary Text **S1**). As an order-of-magnitude estimate, *ϵ* ≃ 3.4 *k*_B_*T* corresponds to the Poisson-Boltzmann energy of a point charge located at 0.4 nm above a surface with uniform charge density comparable to mica, suggesting that electrostatics might indeed affect the dynamics (see Supplementary Text **S1**). A likely more prominent role is, however, expected to be played by internal friction emerging from the intra- and inter-alpha-helices interactions within the coiled coils. Given SbcCD has a low surface charge density (– 0.025 e/nm²), its interaction with the mica surface would be less important than the contacts within the coiled coils. Previous experiments and molecular dynamics simulations of folded proteins revealed subdiffusive dynamics and internal protein friction, and equilibration has been suggested to occur on a second timescale (58–61). These works agree with our results, suggesting that the slow subdiffusive motion reflects the intrinsic dynamics of the coiled coil and is governed by internal friction. Our data do not allow us to estimate the relative importance of mica-protein interaction and internal friction; hence, for further insights we relied on molecular dynamics simulations, as discussed below.

Besides the predominant *S* conformation, we also identified complexes presenting a C-shaped conformation (*C*_*SbcCD*_, ∼8 %, **Fig. 2A-i**); a braided conformation, where the dimeric coiled coil folded at the hook region, and one monomeric segment crossed over the other (*braided*_*SbcCD*_, ∼4 %, **Fig. 2B-i**), producing a rod-like geometry (Supplementary Figure **S8**); and a closed conformation, with the two head domains bound together forming a ring (*closed*_*SbcCD*_, ∼6 %, **Fig. 2C-i**). Remarkably, the contour lengths of these conformations were significantly longer (∼49–55 nm) than that of *S*_*SbcCD*_ (∼44 nm). The *closed*_*SbcCD*_ conformation has been reported before as a result of ATP/ATP-γ-S binding to the MR complex (20, 23). Indeed, the presence of ATP led to a relatively higher probability of observing the closed formation 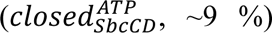 (Supplementary Figure **S9**). Both *braided*_*SbcCD*_ and *closed*_*SbcCD*_ are physiologically relevant for the DNA double-strand break (DSB) repair mechanism. The closed geometry might play a crucial role in effectively encircling DNA during the DNA sensing process, whereas the rod-shaped braided geometry is likely the conformation adopted when enveloping DNA in the DNA-cutting state (20). Interestingly, complexes stochastically and reversibly changed between conformations, mainly from *S*_*SbcCD*_ to *C*_*SbcCD*_ and vice versa, and explored undefined intermediate states, revealing their inherent flexibility and dynamics and a broad and rough energy landscape. The longer contour length and right-handedness of the coiled coils suggest twisting of one head domain and subsequent uncoiling of the coiled coil during the *S*_*SbcCD*_ to *C*_*SbcCD*_ transition (Supplementary Text **S2** and Supplementary Figure **S10**).

**Figure 2:**
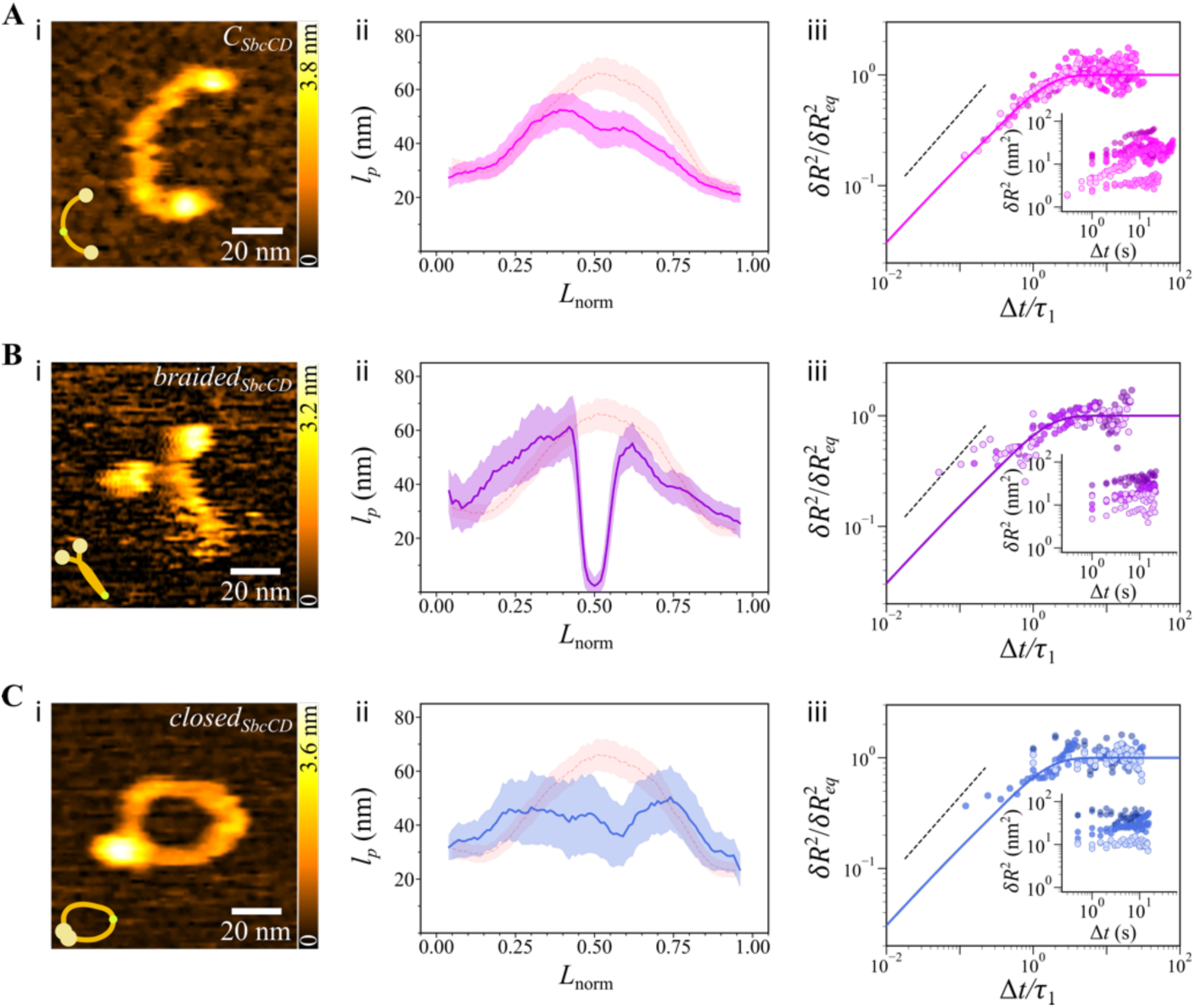
Representative HS-AFM topographs of the different conformations observed for the SbcCD complex with the schematic representation (inset) for *C* (**A-i**), *braided* (**B-i**), and *closed* conformations (**C-i**). Local persistence length *l*_*p*_ as a function of normalized contour length *L*_*norm*_ for the corresponding conformations: *C* (**A-ii**), *braided* (**B-ii**), and *closed* (**C-ii**). The *l*_*p*_ of *S* conformation is shown in the background for comparison. Rescaled MSD as a function of the rescaled time for the respective conformations: *C* (**A-iii**), *braided* (**B-iii**), and *closed* (**C-iii**). The raw MSD data are shown in the insets.

In contrast to *S*_*SbcCD*_, *C*_*SbcCD*_ and *closed*_*SbcCD*_ showed a more uniform flexibility (*l*_*p*_) along the coiled coil length with a more flexible region near and around the hook (**Fig. 2A-ii, B-ii, C-ii**). The sharp decrease in *l*_*p*_ at the hook region of *braided*_*SbcCD*_, falling below the rigidity observed at the globular domain ends region, was possibly an artifact due to the imposition of a continuous arc across the two segments during skeletonization. However, it may reflect a different conformational state of the hook, perhaps switching from an antiparallel to a parallel configuration (62, 63). The observed longer contour length suggests that the conformational state of the complex (*S*_*SbcCD*_, *C*_*SbcCD*_, etc.) is reflected in the coiled coil, possibly through uncoiling, which explains the decreased rigidity of *C*_*SbcCD*_ and *closed*_*SbcCD*_. In contrast, uncoiling of the *braided*_*SbcCD*_ may induce further contacts between the two coiled coils, effectively increasing its *l*_*p*_. Overall, our results suggest that the conformational state of the complex and the interaction between the different molecular components induce changes in the flexibility of the coiled coil region.

To explore the effect of conformation on the dynamics, we determined the MSD over elapsed time as described for *S*_*SbcCD*_. In all conformations, we observed similar behavior, with the MSD scaling as a power law at short times and saturating at longer times (**Fig. 2A-iii, B-iii, C-iii**). Compared to *S*_*SbcCD*_, all three conformations tended to saturate at shorter times (1– 3 s). As suggested by the longer contour length, possible uncoiling may result in lower inter-coil contacts, effectively decreasing internal friction and, thus, the friction coefficient and the relaxation time (**Table 1**). Finally, the dynamic response of the monomeric form (i.e., *monomer*_*SbcCD*_) appeared to be more flexible than the dimeric conformations, while being more dynamic (shorter relaxation time ∼0.8 s, Supplementary Figure **S4** and **Table 1**). This further reveals the role of conformation and multimerization in the flexibility and dynamics of coiled coils. Remarkably, as for *S*_*SbcCD*_, the rescaled *δR*^2^(Δ*t*) for all these conformations collapsed into a master curve (**Eq. 1**). Overall, our observations revealed what we termed an *allodynamic* mechanism. In analogy to allostery, and in addition to it, the molecular composition and conformational state of MR complexes modulate the dynamics and structural flexibility of the coiled coils, modulating the relaxation time and likely affecting biological function.

To determine whether the observed dynamic flexibility was unique to bacterial coiled coils or conserved across species and to assess the possible effect of disease-related mutations, we investigated the eukaryotic homolog hMRN complex (**Fig. 3A**). HS-AFM imaging of the hMRN complex revealed mostly a closed conformation (*closed*_ℎ*MRN*_, **Fig. 3C-i**), resembling *closed*_*SbcCD*_, with coiled coil segments connected at both ends through the hook domain and NBD globular domains. Further, smaller, loosely attached domains were observed to interact dynamically with the globular assembly. This configuration confirmed that the hMRN existed as a heterotetramer on the surface, with two RAD50 monomers forming a dimer through their hook and globular domains, while MRE11 dimers were associated near the globular assembly. Discerning the NBS1 in HS-AFM topographs was challenging due to its small size and mobility. The *closed*_ℎ*MRN*_ conformation was the dominant structure (∼87 %), the rest being open (i.e., *open*_ℎ*MRN*_, **Fig. 3B**). This contrasts with the less abundant closed conformation in SbcCD (*closed*_*SbcCD*_, ∼6 %) and reflects a more stable interaction between the head domains in hMRN. This difference could be a result of a seemingly increased stability of the NBS1-wrapped MRE11 dimer compared to a less stable SbcD dimer, as judged by cryo-EM classes and gel filtration (64, 65). In contrast to *closed*_*SbcCD*_, local determination of *l*_*p*_ of *closed*_ℎ*MRN*_ revealed higher flexibility near the hook, being more rigid at the central regions of each coiled coil segment (**Fig. 3C-ii**). As for SbcCD, the MSD of hMRN increased as a power law at short times but saturated on average at longer times (∼6 s for the *closed*_ℎ*MRN*_, and ∼15 s for *open*_ℎ*MRN*_). Like for SbcCD, the rescaled MSD data of hMRN collapsed into a master curve (**Fig. 3B-iii, C-iii**).

**Figure 3:**
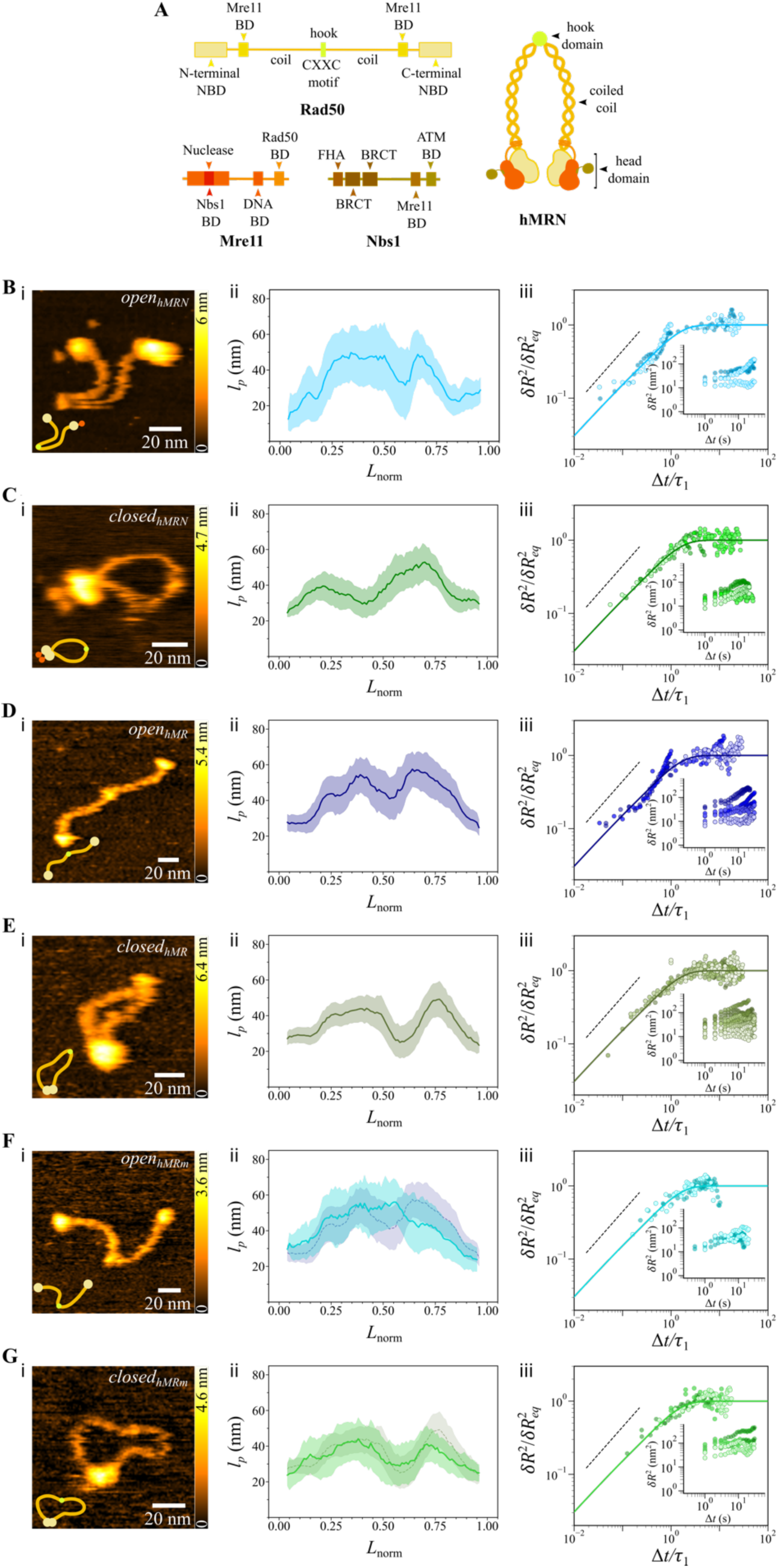
**A.** Left: Scheme of the domain organization of RAD50, MRE11, and NBS1 forming the hMRN complex. Right: Scheme of a dimer of hMRN heterotrimer (MRN)_2_, a RAD50 dimeric coiled coil in the presence of two MRE11 and two NBS1. Representative HS-AFM topographs showing the different conformations of dimers with schematic: *open* (**B-i**) and *closed* (**C-i**) conformations of the hMRN heterotrimeric complex, *open* (**D-i**) and *closed* (**E-i**) conformations of the hMR heterodimeric complex, and *open* (**F-i**) and *closed* (**G-i**) conformations of the hMRm heterodimeric complex. Plots showing the local persistence length *l*_*p*_ as a function of normalized contour length *L*_*norm*_ for the corresponding conformations: *open* (**B-ii**) and *closed* (**C-ii**) for hMRN, *open* (**D-ii**) and *closed* (**E-ii**) for hMR, and *open* (**F-ii**) and *closed* (**G-ii**) for hMRm. The *l*_*p*_ of hMR conformations is shown in the background of the corresponding hMRm conformation for comparison. Plots of the rescaled MSD as a function of the rescaled time for the respective conformations: *open* (**B-iii**) and *closed* (**C-iii**) for hMRN, *open* (**D-iii**) and *closed* (**E-iii**) for hMR, and *open* (**F-iii**) and *closed* (**G-iii**) for hMRm. The raw MSD data are shown in the insets of each rescaled plot.

To assess the influence of protein composition, we imaged hMR (RAD50 + MRE11) and hR (RAD50 alone) **(Fig. 3 and Table 1)**. hMR exhibited open (∼26%, *open*_ℎ*MR*_) and closed (∼74%, *closed*_ℎ*MR*_) conformations, both being longer (∼57 nm) than hMRN (∼51 nm), suggesting uncoiling. The MSD scaling with time was similar to hMRN, with overall shorter relaxation times (**Table 1**), further suggesting uncoiling. hR alone exhibited only the open conformation (*open*_ℎ*R*_), with relaxation time and persistence length comparable to those of closed hMRN. These results reveal that the molecular composition of the complex induces conformational changes and modifies coiled coil flexibility and dynamics.

A naturally occurring mutation in RAD50, characterized by a single amino acid deletion (RAD50^E1035Δ^) within the heptad repeats of the coiled coil near the globular domain, has been shown to impair its function and cause a syndrome involving developmental defects and severe immunodeficiency (66). To explore the effect of this mutation on the dynamic properties of the coiled coil, we visualized the conformational dynamics of the RAD50^E1035Δ^ mutant, both in complex with MRE11 (hMR^E1035Δ^ or hMRm) and alone (hR^E1035Δ^ or hRm) (see Supplementary Figure **S11**). For the hMRm complex, we again found two predominant conformations (*open*_ℎ*MRm*_, ∼52 % and *closed*_ℎ*MRm*_, ∼48 %), but with different relative abundance than for hMR (*open*_ℎ*MR*_∼26 % and *closed*_ℎ*MR*_∼74 %) (**Fig. 3F-i, G-i**). Both hMRm conformations exhibited larger fluctuations in flexibility along the contour length (**Fig. 3F-ii, 3G-ii**), with the closed conformation slightly more flexible. Notably, analysis of the MSD over time revealed shorter relaxation times of the E1035Δ open and closed conformations (∼3 s) than those of the wild type (∼10 s and ∼4 s, respectively) (**Fig. 3F-iii, G-iii** and **Table 1**). Finally, in contrast to wild type hR alone, which only appeared in the open conformation, hRm exhibited both open (*open*_ℎ*Rm*_) and closed (*closed*_ℎ*Rm*_) conformations with longer relaxation times (**Table 1**). Overall, our results reveal that a point deletion in the coiled coil led to different relaxation times and a changed distribution of open/closed conformations, which may partially explain the impaired function that results in the pathogenic state (66).

It is important to highlight that all rescaled MSD across species and conformations collapsed onto a single master curve (**Fig. 4**), following a power law with an exponent of 3/4 at short times and reaching an equilibrium plateau at times longer than the fundamental relaxation time. This observation suggests a universal scaling response conserved across species and well described as semiflexible filaments with strong internal friction.

**Figure 4:**
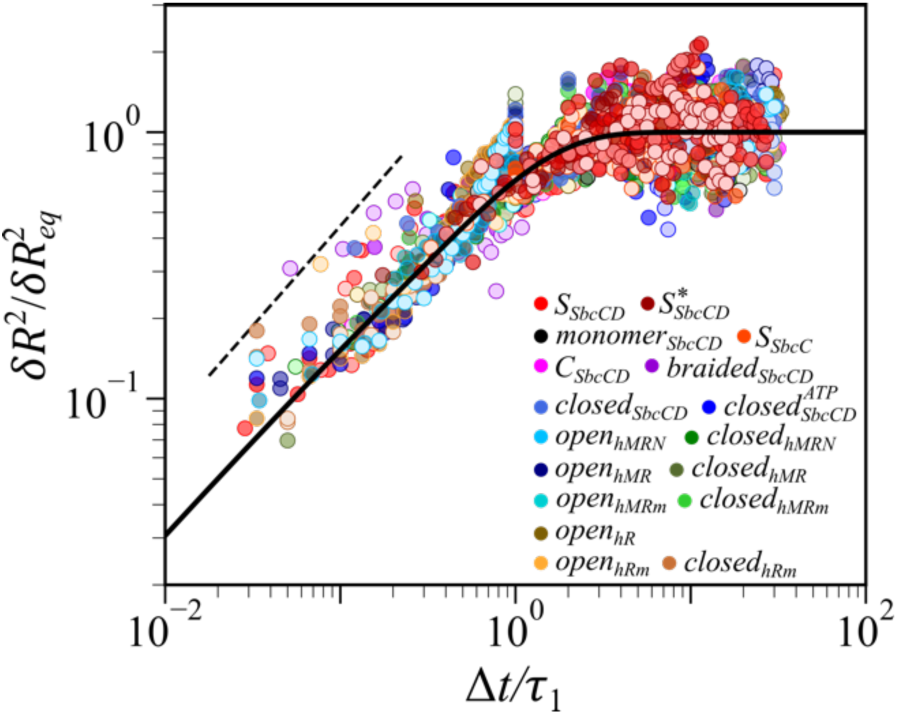
A plot of rescaled MSD in R as a function of the rescaled time for all the conformations of SbcCD, SbcC, hMRN, hMR, hMR^E1035Δ^, hR, and hR^E1035Δ^. The black line represents **Eq. 1**, following a slope of 3/4 (black dotted line) before reaching saturation.

To investigate the molecular origin of this dynamic response, we carried out molecular dynamics (MD) simulations. For that, we extracted the residues of a single coiled coil segment from the SbcC subunit (171 to 904, UniProt code: P13458) and generated the initial structure employing AlphaFold3 (67). We then performed 1 µs-long all-atom MD simulations of this model in an aqueous environment on an ideal, frictionless surface using a confining potential (see Methods). The side view (**Fig. 5A-i**) highlights that the confining potential acts only in the z-direction, allowing the coiled coil to relax freely within the plane and adopt conformations such as in **Fig. 5A-ii**, thereby mimicking the experimental conditions, yet within an ideal two-dimensional confinement.

**Figure 5.**
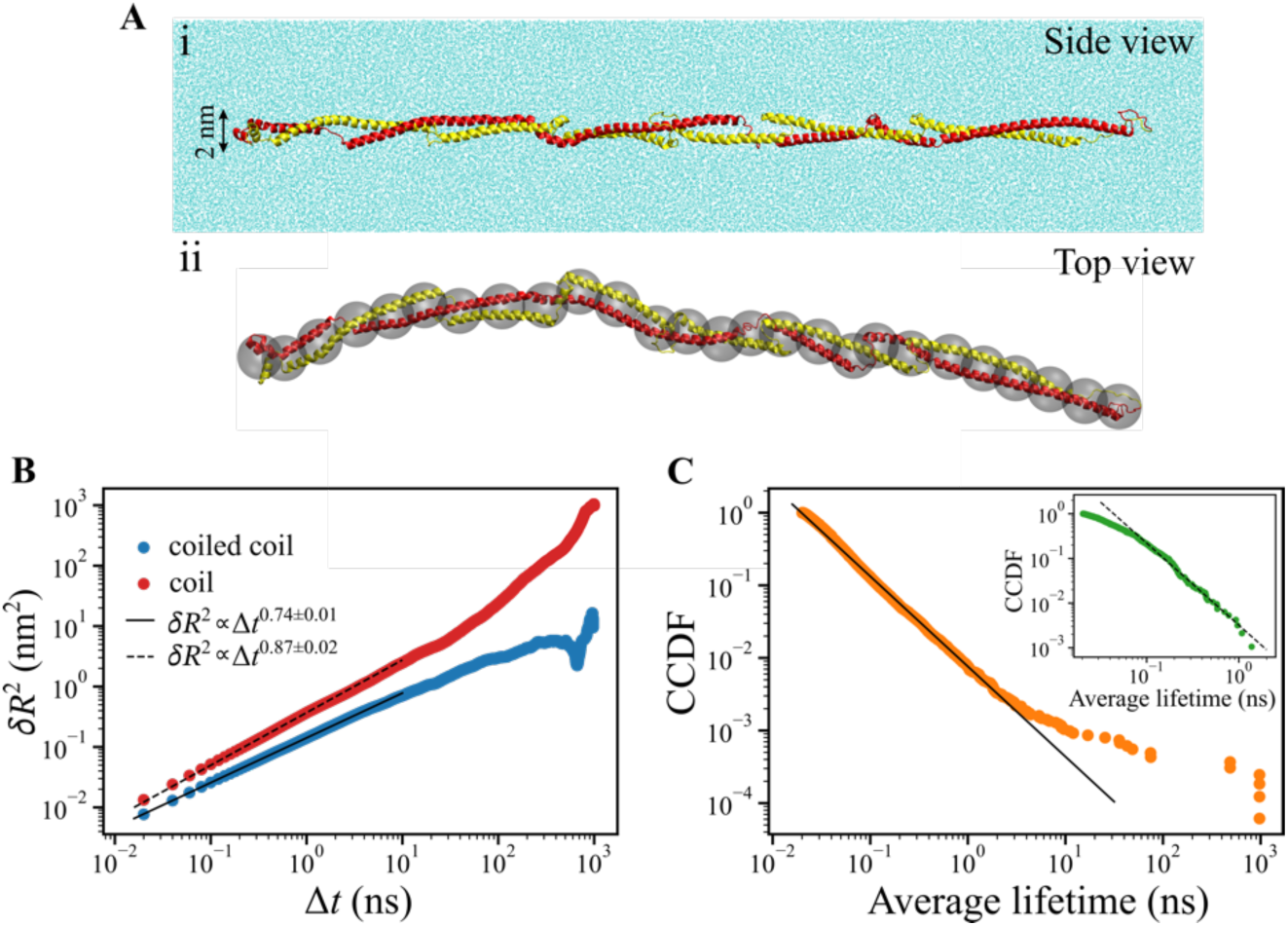
All-atom molecular dynamics simulations. **A**. **i**. Side view of a representative snapshot showing the two-dimensional confinement of the coiled coil. The two coils are represented in red and yellow, while water molecules are depicted as cyan points. To improve visibility, water molecules lying in front of the coiled coil were omitted from the representation. The confining potential restricts the coiled coil to a ∼2 nm slab in the z direction. **ii**. Top view of the same snapshot as in **A**, where water molecules have been omitted to improve visualization. The coarse-grained mapping employed for the analysis of the end-to-end distance is indicated by transparent consecutive grey beads separated by roughly 2 nm. **B**. MSD of the end-to-end distance ⟨*δR*^2^⟩ as a function of elapsed time Δ*t*, calculated from the trajectory of a coiled coil (red circles) or a single coil (blue circles). Data were binned with a bin size equal to 0.05 ns (hydrogen bonds) and 0.67 ns (van der Waals contacts). The black lines are fits to power laws as indicated in the legend. **C**. Complementary cumulative distribution function (CCDF) of average lifetimes of inter-coil contacts detected in the simulation of a coiled coil for van der Waals contacts (main plot) and hydrogen bonds (inset). The black lines are fits to power laws with exponents of −1.2 (solid line) and 1.8 (dashed line).

To assess the end-to-end distance dynamics (**Eq. 1**), each conformation was mapped onto a coarse-grained polymer (**Fig. 5A-ii**, see Methods and Supplementary Information), and the corresponding MSD of the end-to-end distance, *δR*^2^(Δ*t*) was computed. The short-time behaviour of *δR*^2^(Δ*t*) closely follows a power-law function for four orders of magnitude, *δR*^2^(Δ*t*) = *a* · Δ*t*^*b*^, with the exponent *b* = 0.74 ± 0.01 (**Fig. 5B**), in full agreement with the value of 3/4 observed experimentally. From the fitted prefactor *a*, the longest relaxation time in **Eq. 1** is 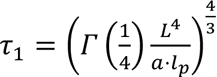 (Supplementary Text **S3**). To determine its value, we computed the contour length *L* = 47 nm (with negligible error) and the persistence length *l*_*p*_ = 44 ± 2 nm, following a procedure akin to the experimental local flexibility analysis (Supplementary Figure **S12**). Plugging these values, we obtain *τ*_1_ = 1.3 ± 0.2 µs. Considering that the TIP3P model employed in the simulations underestimates water viscosity (68), *η*_*TIP*3*P*_ = 0.321 mPa·s, the expectation from theory would be 0.12 µs; hence, the time equilibration from simulation exceeds the prediction by a factor ∼11. Consistent with our results, analysis of a previous simulation (69) of an ideal coiled coil under fast pulling yields an effective drag coefficient ten times larger than expected from theory (Supplementary Text **S4**).

Both the 3/4 power-law exponent and the emergence of internal friction align with our experimental observations. However, our results remarkably depart from theoretical polymer physics predictions. Internal friction in polymeric systems has been described for chromosomes (70) and DNA (71) to be associated with the bending modes. Given that the internal friction in folded proteins arises from local torsional rearrangements (72), this model is expected to be appropriate here, since local torsion is associated with large-scale bending (73). However, when internal friction dominates, as seems to be the case here, this model predicts a power-law exponent shifting from 3/4 to 1 (Supplementary Text **S5**). This is in contrast with our experimental and simulation results, which show dominant internal friction yet maintain the 3/4 exponent. This discrepancy suggests a different mechanism in our system, with an internal friction acting globally (Supplementary Text **S5**). We speculated that the formation and breaking of inter-coil contacts might provide a global mechanism, for which the internal friction would be dominated by relative sliding between the two coils. To test this hypothesis, we reasoned that an isolated coil without such sliding should display an exponent of 1, reflecting bending-dominated internal friction. We thus simulated a single coil under the same confinement (residues 171 to 515, depicted in red in **Fig. 5A**) and confirmed that indeed the autocorrelation function *δR*^2^(Δ*t*) shows an exponent closer to 1 (0.87, **Fig. 5B**). This result strongly supports our ansatz that subdiffusive response and internal friction in the coiled coil arises from the interaction between the two coils, which stiffens while slows down the system. Theoretical models of semiflexible filament bundles may better explain this behaviour (74).

Next, we considered the quantitative discrepancy between the 11-fold increase in equilibration time obtained from the simulations and the 5–7 orders of magnitude observed experimentally. As discussed above, an effect from the interaction with mica in experiments is expected, yet part of this difference may also stem from the different timescales considered. Previous simulations over 13 decades in time revealed self-similar dynamics in a folded protein, with characteristic relaxation times depending on the observation time window (58); these ageing effects might also be present in our system. This view is supported by the inter-coil hydrogen bonds and van der Waals contacts along our 1-µs trajectory of a coiled coil. As shown in **Fig. 5C**, the maximum contact lifetime, i.e., the largest streak of consecutive snapshots with a contact, displays a power law, indicating a highly heterogeneous, long-tailed distribution of lifetimes. We observed hydrogen bonds with maximum lifetimes from tens of picoseconds to about 1 ns (**Fig. 5C** inset). More strikingly, van der Waals contacts spanned from tens of picoseconds up to the microsecond scale, covering all timescales accessible in our simulation. These results strongly suggest that longer-lived contacts should be detected for significantly larger simulations, resulting in a large-timescale slowdown of the dynamics partly covering the gap with the experimental values.

## Discussion

In this work, we proposed an approach based on single-molecule HS-AFM imaging to explore the local flexibility and global dynamics of coiled coils in MR complexes, also applicable to other coiled coil systems. We observed that both flexibility, as determined by the local *l*_*p*_, and dynamics, as quantified by the relaxation time, depend on the conformational state and MR variant. The persistence length varied along the contour length of the coiled coils, revealing the highest variability near the hook region across the different conformations. These changes in the local *l*_*p*_ and global dynamics of the coiled coils likely originated from un- or super-coiling, induced by twists of the globular domains as the complexes adopted different conformations. Our results indicate that different conformations, stoichiometry, and ATP trigger allosteric changes transmitted across the coiled coils that, in turn, modulate their relaxation time.

One of the most notable outcomes of our work is the observation of a universal scaling of the dynamics of all coiled coils studied. Whether this dynamic response is specific to MR complexes or general to any coiled coil domain is still an open question, but one that we can now experimentally tackle. Also, our data analysis approach is partially applicable to other techniques like cryoEM (for flexibility) and smFRET (for dynamics). Our results suggest that at short timescales, the coiled coil’s motion is ruled by its length and persistence length, and is importantly slowed down by internal friction arising from relative sliding between the coils, leading to seconds-long relaxation times, at which the molecule equilibrates. This relaxation time, orders of magnitude longer than expected by pure hydrodynamic drag, is governed by internal friction and tuned by the coiled coil contacts. Of relevance, changes in the structural conformation of the complexes also modulated the dynamic response, leading to shorter or longer relaxation times.

We propose an *allodynamic* mechanism in which the relaxation time of the coiled coils is modulated by the composition and conformational state of the MR complex. Tuning of coiled-coil dynamics and flexibility through changes in the head domains may be necessary for proper function during DSB repair. This leads us to speculate that coiled coils have evolved dynamic properties that tune the relaxation time to carry out function. For example, to efficiently find the DNA damage site or to maintain structural stability once the DNA site has been found. This *allodynamic* mechanism may partially explain the reduced DNA repair capacity of this E1035 variant, which, compared to the normal hMR, exhibited different conformational populations and faster dynamics. Our results also allow us to speculate that mechanical constraints applied to the coiled coil, for example, DNA binding, will lead to different scaling dynamics, as expected for semiflexible filaments under tension (75). This may be relevant to better understand other protein systems presenting coiled coils and in which tension is related to function, like myosins and fibrinogen. The current revolution in structural biology driven by prediction algorithms calls for quantitative tools and theories allowing the prediction of protein conformational dynamics. We anticipate that our approach will facilitate the probing of dynamic flexibility in diverse protein systems, uncovering emergent physical phenomena driving biological functions and establishing a foundation for new theories in protein dynamics.

## Methods

### 1. Protein expression and purification

Details of SbcC, SbcD, and SbcCD protein expression and purification can be found in the following article (20). hMRN was expressed and purified as described in the following reference (76). The purification of hMR and hMR ^E1035Δ^ (hMRm) has also been reported previously (66). Human RAD50 (hR) and RAD50 ^E1035Δ^ (hRm) were obtained as N-terminally tagged MBP fusions after overexpression in suspension 239T cells. Cell pellets were lysed with 0.5 M NaCl, 20 mM Tris-HCl pH 7.5, 1 mM EDTA, 1 mM DTT, 10% glycerol, 0.2% NP40, and supplemented with protease inhibitors (Pierce), clarified by centrifugation at 10,000 x g, and slowly injected on the MPB trap column (Cytiva). The column was extensively washed with 0.5 M NaCl, 20 mM Tris-HCl pH 7.5, 1 mM EDTA, 1 mM DTT, 10% glycerol, and the fusion proteins were eluted in the same buffer supplemented with 20 mM maltose. Proteins were flash frozen in liquid nitrogen and stored at −80 °C.

### 2. AFM sample preparation

A freshly cleaved muscovite mica disc (diameter: 1.5 mm; thickness: 0.1 mm; JBG-Metafix, Montdidier, France) was used as a surface without any further treatment. The mica surface was glued to a glass rod (diameter: 1.5 mm, height: 2 mm), and the assembly was then attached to the sample scanner (on the top of a Z piezo) for imaging. Protein stored in 20 mM Tris (pH 7.5), 125 mM NaCl, and 10% glycerol was thawed, and an intermediate dilution of 80 nM was prepared in an imaging buffer consisting of 25 mM Tris (pH 7.4), 50 mM KCl, 1 mM DTT, 5 mM MgCl_2_, and 1 mM MnCl_2_. A 2 µL aliquot of protein solution, with a concentration ranging from 1.6 to 5 nM, was deposited onto the mica surface and incubated for 5 minutes under a humid hood to allow for partial immobilization of the proteins. Following incubation, the surface was washed several times with the imaging buffer to remove any unbound proteins. HS-AFM imaging was then conducted in the same buffer environment, with ∼100 µL of buffer in the imaging chamber. For experiments involving ATP, it was added directly to the imaging chamber at a final concentration of 0.5–1 mM.

### 3. AFM data acquisition

Imaging was performed in amplitude modulation mode on an SS-NEX HS-AFM (RIBM, Japan) equipped with a standard scanner. We used ultrashort cantilevers with a resonance frequency of ∼600 kHz in liquid, and a nominal spring constant of 0.15 N/m (USC-F1.2-k0.15, NanoWorld, Neuchâtel, Switzerland). During scanning, the free oscillation amplitude of the cantilever was set to 2–3 nm, and the set point for feedback control was kept to 80–90 % of the free oscillation amplitude. Images were acquired at a scan rate of 1–4 frames per second. Typically, the size of a pixel was 0.5–1 nm, while the typical size of a scan area was 300 × 300 nm^2^. All the measurements were performed at room temperature (23–25 °C) in the imaging buffer mentioned above. At least a couple of independent experiments were performed for all the bacterial and human homologs under similar experimental conditions.

### 4. Image processing and analysis

All HS-AFM videos were processed using an in-house developed macro implemented in Fiji (ImageJ) image processing software (https://github.com/centuri-engineering/ProtruDe/) (77). This macro enables the efficient handling of tens of HS-AFM videos containing thousands of images by automating the processing of raw videos to enhance video quality. The image processing workflow mainly involves tilt removal via plane subtraction, flattening to remove large-scale artifacts, and high-frequency noise reduction using a Gaussian filter. In some cases, image smoothening was performed using an in-house software routine developed in MATLAB (MathWorks) available at https://github.com/arin83/U1067/.

To obtain the interpolated *x*-*y* coordinates of the coiled coil trajectories, a multipoint line with B-spline fit was traced along the contour of the coiled coil (excluding the head domains at both ends) using the ImageJ plugin Kappa (48). For the analysis of globular domain volumes using ImageJ, the raw images were converted to binary 32-bit images, and a threshold of pixel intensity was applied to ensure that only the globular domains were considered, excluding the coiled coil regions. The Analyze Particles function was then employed to measure the area and mean grey value of the two globular domains within a dimeric protein complex across a stack of images. These measurements were subsequently used to calculate the volumes of the globular domains.

### 5. Data analysis of coiled coil trajectories

For the assessment of local flexibility and dynamics of each conformation/species, we analysed at least 2 HS-AFM videos (Supplementary **Table S1**). These analyses were performed over all conformations, using data from independent experiments with similar physiological conditions. The descriptions of all the videos can be found in the Supplementary **Table S2**.

### 6. Local flexibility

To extract the persistence length as a function of the coiled coil length, we developed the following routine for each frame of an HS-AFM video. It involved alignment of each molecule, normalization of the contour length, computation of the persistence length through a 2D WLC fit, and averaging.

To align the coiled coil skeletal coordinates across all frames in each video, the skeletal coordinates in each frame were rotated such that the endpoints, corresponding to both globular domains, were positioned along the *x*-axis. This linear transformation of coordinates is distance preserving and results in a centered coordinate system (*x*_*c*_, *y*_*c*_) such that one of the globular domains in all frames is aligned/centred at (0,0).

The contour length (*L*) was calculated along the skeletal coordinates, and then it was normalized (*L*_*norm*_) by dividing by the protein contour length in that frame, ensuring *L*_*norm*_ ranged between [0,1]. For all analyses in this work, the midpoint (corresponding to the hook domain) of the dimeric coiled coil was considered to be at an *L*_*norm*_ of 0.5, with the globular domains of each monomeric segment at *L*_*norm*_ of 0 and 1, respectively. This provides the coordinate range for one monomeric segment as *L*_*norm*_ = (0, 0.5) and for the other as (1, 0.5).

To compute the local persistence length (*l*_*p*_), a rolling window approach was employed. A constant window size of 10 nm was set and rolled along the entire contour length of the dimeric coiled coil in increments of 1 nm, from one end to the other. Within each window, the skeletal coordinates of the coiled coil were partitioned into 10 equal segments. For each segment, the squared end-to-end distance (*R*^2^) and the corresponding arc length (*L*_*arc*_) were calculated. These data points were then fitted using a 2D worm-like chain (WLC) model to extract the local *l*_*p*_ for each window position, as illustrated in **Fig. 1D** (49):

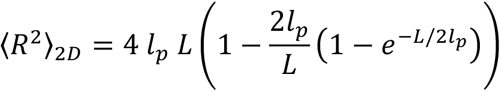

This allowed us to obtain a value of *l*_*p*_ at each point along the normalized length *L*_*norm*_. For every conformation, the local *l*_*p*_ at each window was binned and averaged across all frames from all the HS-AFM videos analyzed.

### 7. Dynamics of coiled coil segments

To study the dynamics of the various conformations and molecular composition of SbcCD and hMRN (including mutants) complexes, we focused our fluctuation analysis on two key metrics: the time trajectories of the coiled coil’s end-end distance (*R*) and the angle between the two ends of the coiled coil (*θ*_*mid*_). For each coiled coil segment within each frame of a HS-AFM video, *R* was defined as the distance between the coiled coil’s globular domain and the hook domain. Subsequently, *θ*_*mid*_ was defined as the angle between two vectors, one extending from the middle point of the segment to the hook and the second from that same middle point to the head domain (Supplementary Figure **S7**). Using the scan rate of each HS-AFM video, we determined the time (in seconds) from the frame number.

We then computed the mean square deviation (MSD) in both *R* and *θ*_*mid*_ as a function of the elapsed time Δ*t* as

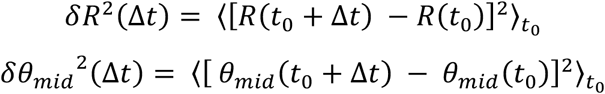

where *t*_0_ denotes the initial time, and the mean was calculated over all possible values of *t*_0_. To ensure a robust statistical description of the differences and an adequate sample size for calculating the mean at longer Δ*t*, the MSD was calculated until a maximum elapsed time of half the total number of frames in each HS-AFM video.

To determine the effective persistence length, internal friction coefficient, and relaxation time from the dynamics of each coiled coil segment, the following expression was fitted to the *δR*^2^(Δ*t*) (78):

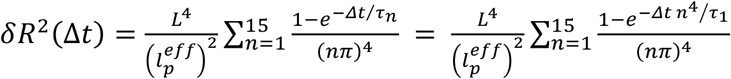

where the relaxation time of the fundamental mode 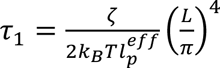, the relaxation time of mode 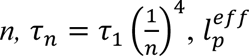 is the effective persistence length, *ζ* is the internal friction coefficient, *k*_B_ is the Boltzmann constant, and *T* is the temperature. For the non-linear fitting, the first 15 terms of the above series were considered, and the fitter parameters were 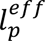 and *τ*_1_, using the time-averaged contour length (*L*) from each video. The fitting bounds were set to 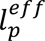 ∈ [0.1, 1000] and *τ*_1_ ∈ [0, max(Δ*t*)], and the SEM was used for weighting in χ^2^ minimization.

Within each HS-AFM video, the *δR*^2^(Δ*t*) of each monomeric segment was rescaled by its characteristic equilibrium saturation value 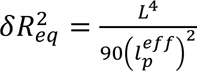 and the elapsed time by its characteristic (fundamental) relaxation time *τ*_1_ (52).

### 8. Code availability

All the analysis procedures presented in sections 6 and 7 were performed using custom-written modular Python code. The source code is freely available on GitHub (source code: https://github.com/DyNaMo-INSERM/PyDycoil).

### 9. Molecular dynamics simulations

All-atom molecular dynamics were performed on a single coiled coil extracted from the SbcCD protein complex. We provided the full sequence of the coiled coil as input to AlphaFold3. From the best-scoring structure, we extracted residues 171 to 904, which provided the initial coordinates for our simulations of the coiled coil. For a single coil, we restricted the system to residues 171 to 503 (note that the red coil in **Fig. 5** corresponds to residues 171 to 515; for the single coil system, we discarded the 12 disordered residues in the region of the zinc hook).

From the initial coordinates, the simulation box was built with the program tLeap from AmberTools (79), by considering the ff14SB force field for the protein (80) and the TIP3P model for water (81). To ensure overall electroneutrality, we added a suitable number of sodium ions, parameterized according to the Joung-Cheatham prescription (82). We subsequently converted the topology and coordinate files produced by tLeap to the GROMACS format by means of ParmEd, also belonging to AmberTools (79).

Simulations were then performed in GROMACS 2025.2 (83). We first minimized the system by performing up to 5×10^4^ steepest descent steps. Then, we introduced a flat-bottomed potential restraining the protein onto a planar slab of 2 nm thickness parallel to the x-y plane via a harmonic energetic penalty with spring constant 100 kJ/(mol·nm^2^). This protocol allowed mimicking adsorption to a substrate; however, the protein was still completely surrounded by water, which in practice corresponds to an ideal, frictionless adsorbing surface. With this setup, we proceeded to a 20-ns simulation at constant temperature (300 K) and pressure (1 bar), to allow relaxation of the box volume. Temperature was maintained via a Langevin thermostat with a 2 ps relaxation time, while the barostat was implemented via C-rescale (84). Finally, we proceeded to 1 *μ*s NVT production simulations in the presence of the flat-bottomed potential. In all our simulations, we imposed a cutoff of 1.2 nm for the van der Waals and electrostatic interactions; long-range electrostatics was implemented via particle-mesh Ewald. For the NPT and NVT simulations, we set an integration timestep equal to 2 fs, enabled by constraining the bonds involving hydrogens via the LINCS algorithm. In the production phase, we saved the coordinates every 20 ps.

To study the polymer physics of the coiled coil, we mapped the atomic coordinates onto a coarse-grained representation, where effective beads were separated on average by roughly 2 nm (Supplementary **Table S3)**. Starting from the coarse-grained representation, we computed the contour length of the coiled coil (or single coil) by summing the distances between consecutive effective beads, while the end-to-end distance was defined based on the separation between the first and last beads.

We computed hydrogen bonds and van der Waals contacts based on geometric criteria. We considered a hydrogen bond as formed when the following criteria were satisfied simultaneously (85): i) distance between donor and acceptor below 0.35 nm; ii) distance between hydrogen atom and acceptor below 0.245 nm; iii) angle between donor, hydrogen, and acceptor above 150°; iv) donor and acceptor were either oxygen or nitrogen atoms. We identified van der Waals contacts by considering heavy atoms separated by less than 0.4 nm and with at least four bonds of separation along the topology of the protein. For each inter-coil contact, we assigned a 0 or 1 to each trajectory frame according to whether the contact was detected or not. Then, we computed the average lifetime as the mean length of streaks of consecutive 1s. To mitigate the possibility of false negatives based on mild violations of the geometric criteria, we considered as a single event any two streaks separated by a single frame.

## Acknowledgement

This project has received funding from the Human Frontier Science Program (HFSP, grant no. RGP0056/2018 to FR), the European Research Council (ERC) under the European Union’s Horizon 2020 research and innovation program (grant no. 772257 to FR), the German Research Foundation (DFG HO2489/9-1, HO2489/11-1 and EXC 3113 to KPH), the Institut du Cancer (INCa; grant No. 2019-1-PL BIO-01/2019-123 to MM and PR), the Spanish MCIN/AEI/10.13039/501100011033 and FSE+ through a Ramón y Cajal Fellowship (RYC2022-037744-I to SA), and the Spanish MCIN (PID2023-149150OB-I00 to SA and “María de Maeztu” Programme for Units of Excellence in R&D grant CEX2023-001316-M). We thank Shin Morioka and Mikihiro Shibata for their helpful discussions and experimental assistance during the BioSPM summer school in Kanazawa, Japan. We also thank Lisa Käshammer and Fabian Gut for providing purified SbcCD and Johannes Stigler for fruitful discussions.

## Competing interest statement

The authors declare no competing interests.

## Author contribution

Conception: PS, MM, FR; Supervision: MM, FR; AFM data acquisition: PS, AM, MM; Image analysis: PS; Data analysis: YS, PS, FR; Protein expression and purification: PR, TTP, KPH, MM; Molecular dynamics simulations: SA; Manuscript writing: PS, YS, CV, SA and FR, with contributions from all the authors.

## Notes

### Competing Interest Statement

The authors have declared no competing interest.

