## Supplementary Information for "Scaling the Dynamics of Coiled Coils"

**This PDF file includes:**

Supplementary Texts S1 to S5

Supplementary Figures S1 to S12

Supplementary Tables S1 to S3

References for SI

### Supplementary Texts

#### S1. Estimations of mica-protein and inter-coil interactions

The electrostatic interaction between the SbcCD dimeric complex and the mica surface can be evaluated by comparing their respective charge densities. The bare mica surface carries a net negative charge, with a surface charge density of ≈ 2.08 charges/nm² (area per charge ≈ 0.48 nm²) (1). The SbcC subunit contains 135 negatively charged residues and 113 positively charged residues, yielding a net charge of −22e. The surface area of the SbcC calculated using PyMOL is 880.43 nm², which gives a charge density of ≈ 0.025 charges/nm². The sparse population of charges likely contributes to the ruggedness of the bending free-energy landscape. As an order-of-magnitude estimation, we consider here the simple case of the interaction free-energy between a point unit charge and a uniformly charged surface with density $\sigma \simeq$-2.08 charges/nm², mimicking mica. Based on the Poisson-Boltzmann equation, the electrostatic potential $\psi(z)$ at distance $z$ from the surface is regulated by the Gouy-Chapman formula (2):

$$\psi(z) = -\frac{2k_BT}{e_0}\ln\frac{1-\gamma\, e^{-\frac{z}{l_D}}}{1+\gamma\, e^{-\frac{z}{l_D}}}\,.$$

In the previous formula, $k_BT$ is the thermal energy, $e_0 = 1.6 \cdot 10^{-19}$ C the elementary charge, $l_D$ the Debye length of the buffer, and $-1 \leq \gamma \leq 0$ a coefficient quantifying the strength of the surface charge density as compared to the screening action of the buffer counterions. In our buffer, we estimate the ionic strength to be $I \simeq 90$ mM. Based on this, the Debye length is $l_D = 1/\sqrt{8\pi N_A l_B I} \simeq 1$ nm, where $l_B \simeq 0.7$ nm is the Bjerrum length and $N_A = 6.022 \cdot 10^{23}$ mol$^{-1}$ is Avogadro's number. The coefficient $\gamma$ is then obtained as $\gamma = \frac{l_{GC}}{l_D} - \sqrt{\left(\frac{l_{GC}}{l_D}\right)^2 + 1} \simeq -0.9$, where $l_{GC} = 1/2\pi l_B\sigma \simeq 0.1$ nm is the Gouy-Chapman length. For the electrostatic interaction of a unit-charge particle with the surface, the Gouy-Chapman formula predicts an energy equal to $e_0\psi(z)$, yielding 3.4 $k_BT$ at a distance roughly equal to 0.4 nm. In this picture, the specific position of coiled coil charges along the mica surface introduces a ruggedness on the bending free-energy landscape of the protein, for which this estimation provides an order-of-magnitude estimate.

However, internal friction of the coiled coil is also expected to provide a significant contribution to such ruggedness. As discussed in the main text, we ascribe the main mechanism of internal friction to be due to inter-coil contacts. The relative sliding between the two coils combined with bending provides a heterogeneous set of relative arrangements between the various charges present in the coiled coil, contributing to the free-energy ruggedness in a similar fashion as discussed above for mica. The relative sliding is regulated by inter-coil hydrogen bonds and van der Waals contacts, which, from our MD simulations, were estimated to be established with a linear density of $1.70 \pm 0.01$ and $50.1 \pm 0.1$ contacts/nm, respectively.

As an estimation of the expected relative importance of internal friction as compared to interaction with mica, we assumed that all heavy atoms of the coiled coil found within the bottom 0.4 nm layer of the confining box would establish van der Waals contacts with the mica substrate, yielding a linear density of $24.0 \pm 0.5$ contacts/nm, i.e., roughly half the inter-coil contacts computed above. All in all, this analysis suggests that internal friction is at least an equal player as mica interaction in determining the equilibration timescale observed in experiments, and it likely provides the main contribution.

#### S2. Conformational transition

The dimeric coiled coil of the SbcCD complex displayed distinct local flexibility and global dynamics across different conformations. We attributed this to differences in the degree of coiling of the coiled coil structure. In accordance, the same coiled coil segment may feature higher or lower stiffness and faster or slower dynamics, likely due to super- or under-coiling.

To understand how conformational changes induce mechanical and dynamical changes, we took advantage of the trajectories that show a reversible switch from one conformation to the other. For that, we focused on the relatively frequent $S_{SbcCD}$-to-$C_{SbcCD}$ conformational transition (Supplementary Figure **S10**). The $S_{SbcCD}$ was characterized by the change from negative to positive curvature, and the $C_{SbcCD}$ by a continuously positive curvature. Other conformations found during this transition were considered non-classified conformations. We were unable to track the intermediate states due to the limited time resolution of HS-AFM imaging, but we found that the conformational transition was relatively fast, occurring mostly from one frame to the next, and was reversible and stochastic.

Given the inherent curvature of each monomeric coiled coil segment, we hypothesized that, for the $S_{SbcCD}$-to-$C_{SbcCD}$/$C_{SbcCD}$-to-$S_{SbcCD}$ transition to occur on a surface, one of the globular domain ends in the dimeric coiled coil must turn or twist to change its orientation, from an upward to a downward position. In this out-of-plane transition, the twist of a head domain induces the monomeric coiled coil segment to flip either clockwise or counterclockwise around the longitudinal axis, depending on the direction of the head twist. This rotational motion may induce super-coiling or under-coiling, potentially altering the local flexibility and the dynamics between the two conformations. These deformation-induced changes in local flexibility are not only relevant to the $S_{SbcCD}$-to-$C_{SbcCD}$/$C_{SbcCD}$-to-$S_{SbcCD}$ transition but applies to other conformational states. For an in-depth understanding, it would be important to develop a model that incorporates this out-of-plane rotation while accounting for the electrostatic interactions with the underlying mica surface or to carry out mechanical measurements upon under- and super-coiling using, for example, magnetic tweezers. Given the longer contour length and increased flexibility of the $C_{SbcCD}$ conformation, we propose that its coiled coil is under-coiled as compared to the $S_{SbcCD}$ conformation.

**S3. Short-time behavior of correlation function**

As reported in the main text, the dynamics of the end-to-end distance $R_{2D}$ of a wormlike chain in two dimensions with length $L$ and persistence length $l_p$ is given by

$$\langle \delta R^2 \rangle = \langle [R_{2D}(t+\Delta t) - R_{2D}(t)]^2 \rangle_t = \frac{L^4}{\pi^4 l_p^2} \sum_{n=1}^{\infty} \frac{1 - e^{-\frac{\Delta t}{\tau_1} n^4}}{n^4}, \qquad (1)$$

where

$$\tau_1 = \frac{L^4 \, \zeta}{2 k_B T l_p \pi^4} \qquad (2)$$

is the longest relaxation mode, with $\zeta$ being the transverse drag coefficient per unit length of the filament. We derive here the short-time power-law behavior of $\langle \delta R^2 \rangle$. The key point is to observe that one can write

$$1 - e^{-\frac{\Delta t}{\tau_1} n^4} = e^{-\frac{\Delta t'}{\tau_1} n^4} \Big|_{\Delta t' = \Delta t}^{0} = -\int_0^{\Delta t} d\Delta t' \frac{d}{d\Delta t'} e^{-\frac{\Delta t'}{\tau_1} n^4} = \frac{n^4}{\tau_1} \int_0^{\Delta t} d\Delta t' \, e^{-\frac{\Delta t'}{\tau_1} n^4} .$$

Hence, we obtain

$$\sum_{n=1}^{\infty} \frac{1 - e^{-\frac{\Delta t}{\tau_1} n^4}}{n^4} = \frac{1}{\tau_1} \int_0^{\Delta t} d\Delta t' \sum_{n=1}^{\infty} e^{-\frac{\Delta t'}{\tau_1} n^4} ,$$

where we inverted sum and integral. Since we are considering $\Delta t' \ll \tau_1$, we can approximate $\sum_{n=1}^{\infty} \simeq \int_0^{\infty} dn$, hence:

$$\sum_{n=1}^{\infty} \frac{1 - e^{-\frac{\Delta t}{\tau_1} n^4}}{n^4} \simeq \frac{1}{\tau_1} \int_0^{\Delta t} d\Delta t' \int_0^{\infty} dn \; e^{-\frac{\Delta t'}{\tau_1} n^4} .$$

We define the variable

$$u \equiv \frac{\Delta t' n^4}{\tau_1} \Rightarrow n = \left(\frac{\tau_1}{\Delta t'}\right)^{\frac{1}{4}} u^{\frac{1}{4}} \Rightarrow dn = \frac{1}{4}\left(\frac{\tau_1}{\Delta t'}\right)^{\frac{1}{4}} u^{-\frac{3}{4}} du\,.$$

Hence, we obtain

$$\sum_{n=1}^{\infty} \frac{1 - e^{-\frac{\Delta t}{\tau_1} n^4}}{n^4} \simeq \frac{1}{4\,\tau_1^{\frac{3}{4}}} \int_0^{\Delta t} d\Delta t' \; (\Delta t')^{-\frac{1}{4}} \int_0^{\infty} du \; u^{-\frac{3}{4}} e^{-u} \;.$$

The latter integral is by definition $\Gamma(1/4) \simeq 3.626$, where $\Gamma$ is Euler's Gamma function. The former integral is also easily solved, yielding

$$\sum_{n=1}^{\infty} \frac{1 - e^{-\frac{\Delta t}{\tau_1} n^4}}{n^4} \simeq \frac{\Gamma\left(\frac{1}{4}\right)}{3} \left(\frac{\Delta t}{\tau_1}\right)^{\frac{3}{4}} \,.$$

Substituting into Eq. (1), we finally obtain the short-time correlation function:

$$\langle \delta R^2 \rangle \simeq \frac{\Gamma\left(\frac{1}{4}\right)}{3\pi^4} \frac{L^4}{l_p^2} \left(\frac{\Delta t}{\tau_1}\right)^{\frac{3}{4}} \,. \tag{3}$$

The master curve is obtained by renormalizing the time as $\overline{\Delta t} \equiv \Delta t/\tau_1$ and the squared correlation by means of the plateau at large times, $\langle \delta l^2 \rangle \equiv 90 \langle \delta R^2 \rangle l_p^2 / L^4$:

$$\langle \delta l^2 \rangle \simeq \frac{30\Gamma\left(\frac{1}{4}\right)}{\pi^4} \; \overline{\Delta t}^{\frac{3}{4}} \simeq 1.11661 \; \overline{\Delta t}^{\frac{3}{4}} \tag{4}$$

#### S4. Estimation of effective drag coefficient from previous simulation work

In Ref. (3), the authors combine MD simulations and a continuum approach to study the response of an ideal coiled coil to longitudinal pulling. From their analysis, they obtain an $\zeta_{\parallel} \simeq 8 \cdot 10^{-9}$ pN·s/nm$^2$; however, in their case this quantity corresponds to the effective parallel drag coefficient per unit length. Based purely on solvent friction, one expects $\zeta_{\parallel,0} = 2\pi\eta / \ln(2L/r)$ (4), using $L \simeq 18$ nm as the contour length of the coiled coil considered in the published work, while $r \simeq 1$ nm is the radius of the coiled coil as estimated in the main text of our work. Considering that in Ref. (5) the authors employed the SPC model water, for which $\eta = 0.47$ mPa·s is about half the experimental value, the theoretical prediction for a free filament is $\zeta_{\parallel,0} \simeq 0.8 \cdot 10^{-9}$ pN·s/nm$^2$, i.e., simulations yield a drag coefficient about ten times larger than expected, is in line with our MD simulations.

#### S5. Impact of internal friction on short-time behavior of correlation function

Following the approach from Refs. (6, 7), if one assumes internal friction to be associated with the bending modes of the polymer, Eq. (1) is rewritten as

$$\langle \delta R^2 \rangle = \frac{L^4}{\pi^4 l_p^2} \sum_{n=1}^{\infty} \frac{1 - e^{-\frac{\Delta t}{\tau_n}}}{n^4} \,, \tag{5}$$

where

$$\tau_n = \frac{\tau_1}{n^4} + \tau_{int} \,, \tag{6}$$

with $\tau_1$ defined according to Eq. (1) and

$$\tau_{int} = \frac{r^4\,\zeta_{int}}{k_B T\, l_p}. \tag{7}$$

In Eq. (7), $r$ is the radius of the cross section of the polymer (assumed to be a homogeneous and isotropic rod) while $\zeta_{int}$ quantifies the internal friction.

In our experiments and simulations, the fitting value for the drag coefficient is much larger than the expectation from theory $\zeta_0 = 4\pi\eta/\ln(2L/r)$ (with $\eta$ being solvent viscosity), which indicates a prominent role of internal friction. However, in Eq. (6) this implies $\tau_{int} \gg \tau_1$, that is, $\tau_n \simeq \tau_{int}$ independently of $n$. Within this approximation, and exploiting the known result $\sum_{n=1}^{\infty}\frac{1}{n^4} = \frac{\pi^4}{90}$, Eq. (5) becomes

$$\langle \delta R^2 \rangle \simeq \frac{L^4}{90\, l_p^2}\left(1 - e^{-\frac{\Delta t}{\tau_{int}}}\right),$$

which in the short-time limit $\Delta t \ll \tau_{int}$ yields

$$\langle \delta R^2 \rangle \simeq \frac{L^4}{90\, l_p^2} \cdot \frac{\Delta t}{\tau_{int}} \propto \Delta t\,,$$

i.e. a scaling exponent equal to 1.

The only way to maintain a scaling equal to 3/4 is to consider an internal-friction mechanism acting globally (akin to the effect of an increased solvent viscosity), providing relaxation times

$$\tau_n = \frac{\tau_1}{n^4} + \frac{\tau_{int}}{n^4},$$

which in turn ensures a short-time scaling of the correlation function with exponent equal to 3/4 whilst allowing for a larger effective viscosity.

#### Supplementary Figures

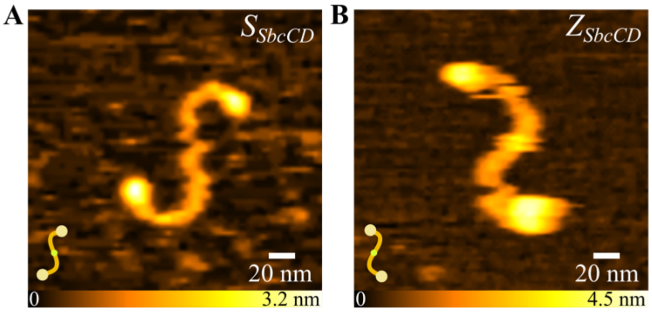


**Supplementary Figure S1. $S_{SbcCD}$ vs $Z_{SbcCD}$ conformations.** Representative HS-AFM images of SbcCD complex deposited on mica reveal predominantly **A**. S-shaped conformations ($S_{SbcCD}$), along with a small fraction (<1 %) of its mirror-symmetric **B.** Z-shaped conformation ($Z_{SbcCD}$). This pronounced asymmetry in conformation distribution (~70 % vs <1 %) indicates a preferred interaction of globular heads with the mica surface while the coiled coil regions remain flexible.

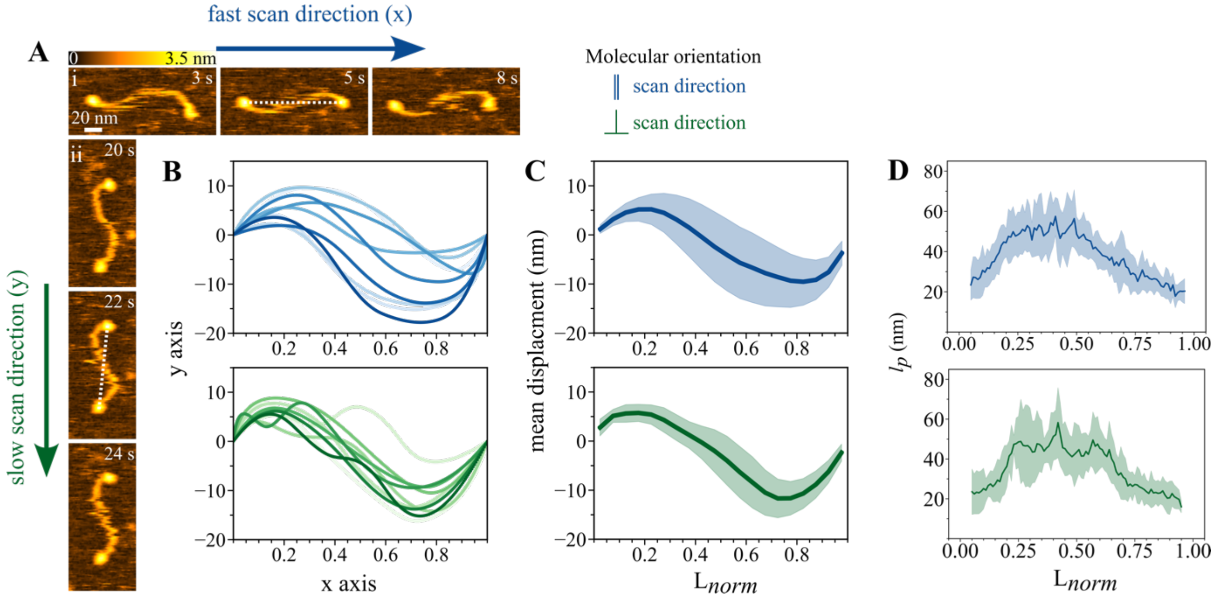


**Supplementary Figure S2. Impact of scan directions on AFM imaging. A.** Sequential AFM images from the same HS-AFM video showing the conformational dynamics of two SbcCD complexes oriented parallel **(i)** and perpendicular **(ii)** to the fast-scanning direction. The displacement of the coiled coil region was observed with respect to their principal molecular axis, as shown with a white dotted line. Timestamps are shown at the top right in each image. **B.** Skeletal coordinates data for multiple frames from the HS-AFM video for both scan directions (top: parallel, and bottom: perpendicular to the fast scan direction). For the analysis, the molecules were aligned with their longitudinal/principal axis parallel to the x-axis (head-to-head dotted line in **A**). **C.** Mean ± standard deviation of the displacement values of the coiled coil region for the two molecular orientations. The middle part of the dimeric structure (near the hook region) shows considerable displacement, indicating minimal HS-AFM tip-assisted bias during scanning. **D.** Plots showing the variation of $l_p$ as a function of $L_{norm}$ for these coiled coil complexes oriented in the fast (top) and slow (bottom) directions of the scan, showing similar flexibility, again suggesting little disturbance of the scanning HS-AFM tip.

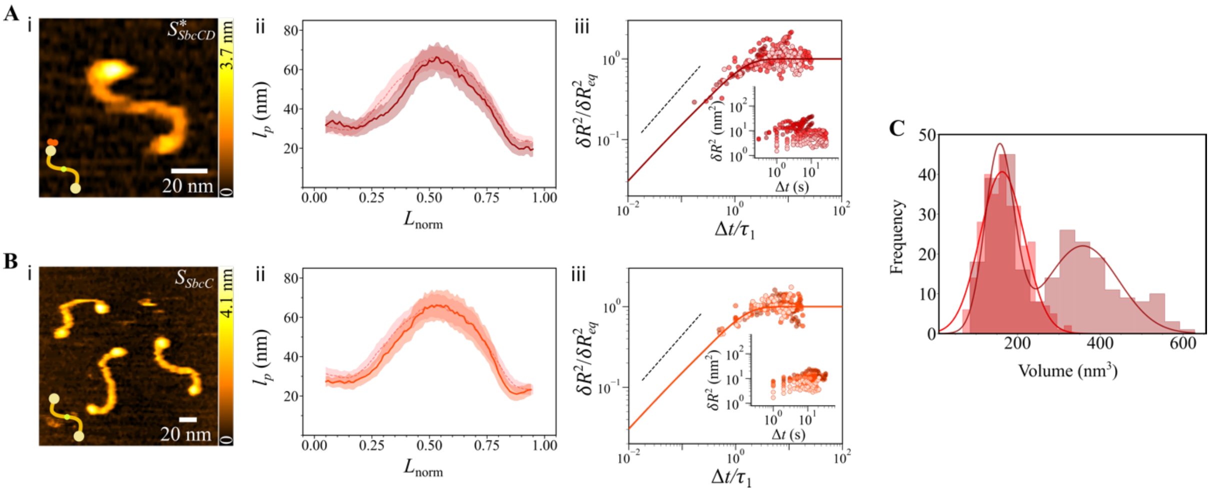


**Supplementary Figure S3. Effect of the presence of SbcD on coiled coil flexibility and dynamics.** **(A-i)** A representative HS-AFM topograph of the $S^{*}_{SbcCD}$ conformation, in which one head presents a larger volume than the other. The presence of SbcD is not discernible in the HS-AFM image, although the two proteins (i.e., SbcC, SbcD) in the SbcCD complex were present at an equimolar ratio during deposition onto the mica surface. **(A-ii)** Local persistence length $l_p$ as a function of normalized contour length ($L_{norm}$) for $S^{*}_{SbcCD}$ with the $l_p$ of $S_{SbcCD}$ in the background. Like for the $S_{SbcCD}$, the central region of the $S^{*}_{SbcCD}$ coiled coil exhibited increased rigidity, whereas regions adjacent to globular domains were more flexible (**Table 1**). **(A-iii)** Rescaled MSD as a function of the rescaled time collapsed into a master curve (dark red solid line). The black dashed line shows a power law with exponent 3/4. Inset: Raw MSD data of $S^{*}_{SbcCD}$. From the fitted parameters, a shorter relaxation time was obtained for the $S^{*}_{SbcCD}$. This suggests that the presence of SbcD may tune the dynamic response of the coiled coil. **(B-i)** HS-AFM imaging of SbcC protein alone under similar experimental conditions predominantly showed an S-shaped conformation (i.e., $S_{SbcC}$, > 95%). The presence of globular ends at both termini confirms that SbcC forms a dimeric coiled coil. No significant volume differences were observed between the two globular ends. **(B-ii)** Plot showing the variation of local persistence length $l_p$ as a function of normalized contour length $L_{norm}$ for $S_{SbcC}$, with the $l_p$ of $S_{SbcCD}$ shown in the background. No significant differences in local flexibility were detected between the two conformations. **(B-iii)** Rescaled MSD as a function of rescaled time collapses onto a master curve (red solid line). The black dashed line shows a power law with exponent 3/4. Inset: Raw MSD data of $S_{SbcC}$. **(C)** A histogram of volume distribution of the globular domains in the $S^{*}_{SbcCD}$ (dark red) shows two distinct populations, while the $S_{SbcC}$ (light red) shows only a single distribution that overlaps with the smaller volume population of the $S^{*}_{SbcCD}$. The populations were fitted to a Gaussian function. For $S^{*}_{SbcCD}$ (sampling size: n= 250), the maxima of the first and second peaks represented the mean volumes of the regular and larger globular domains (156 ± 34 nm$^3$, 357 ± 86 nm$^3$), respectively, with one globular domain ~2 times larger than the other. For $S_{SbcC}$ (sampling size: n= 300), the mean volume of the globular domain was 162 ± 53 nm$^3$. The data are displayed from 5 HS-AFM videos with tens of frames for each of these conformations. Notably, in ~8 % of the population, we observed a volume discrepancy between globular domains at the two ends of the dimeric coiled coil. These findings suggest that the $S^{*}_{SbcCD}$ arises when a SbcD monomer binds to only one of the SbcC NBD termini. In other words, comparison of head domain volume between the SbcC protein alone and the SbcCD complex suggests that the S-shaped SbcCD complex population lacked SbcD in the majority of cases.

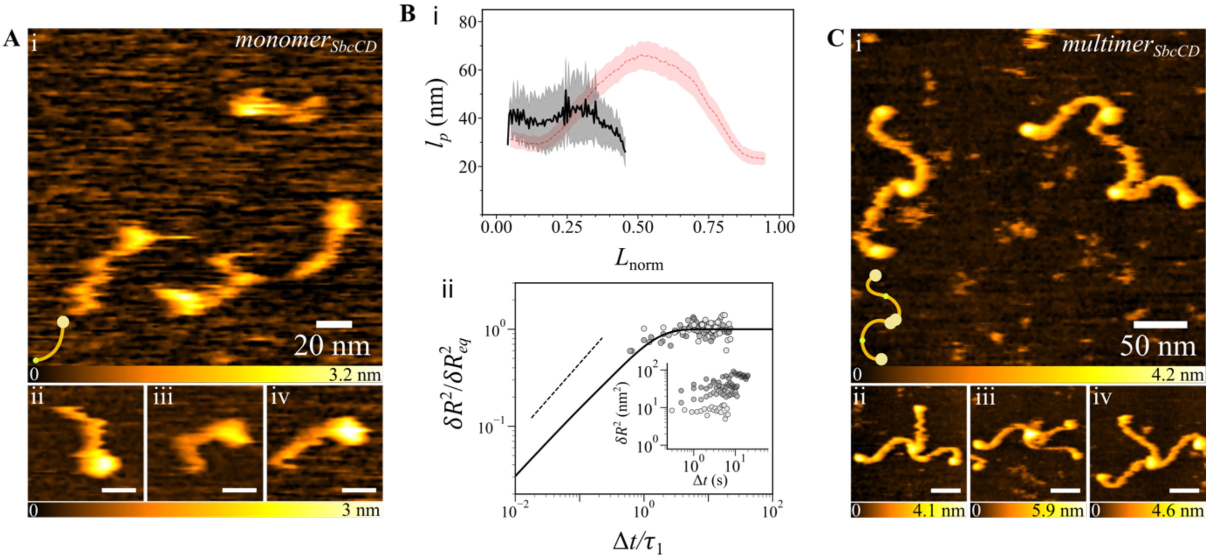


**Supplementary Figure S4. SbcCD in monomeric and multimeric forms. A. (i)** Large area view of SbcCD in the $monomer_{SbcCD}$ coiled coil form and **(ii-iv)** different frames showing its coiled coil's flexibility. In a few instances, the coiled coil exhibits bending near its central region. The scarcity of this conformation on the surface made it difficult to reliably identify and characterize. **B. (i)** To investigate how the presence of a free hook region modulates the mechanical behavior of the coiled coil in this monomeric form, we analysed the local $l_p$ as a function of $L_{norm}$ and compared it to the $l_p$ of $S_{SbcCD}$ as a reference. The data reveal nearly uniform flexibility along the entire contour length, in contrast to the varying flexibility of the dimeric coiled coils. Due to the minimal interaction with the surface, the $l_p$ of $monomer_{SbcCD}$ form was difficult to quantify, as reflected in the larger uncertainty. **(ii)** Rescaled MSD of end-to-end distance as a function of the rescaled time (symbols), which collapsed into a master curve (solid line). Inset: Raw MSD data of $monomer_{SbcCD}$. Shades of gray show different videos. **C. (i)** Large area view of SbcCD multimers where more than two $S_{SbcCD}$ are connected via their globular ends, forming a concatenated chain structure. **(ii-iv)** Different frames show the dynamics of $multimer_{SbcCD}$ conformers where three $S_{SbcCD}$ are associated via globular domains, while their free globular ends move flexibly in different spatial orientations. The abundance of the multimers was increased by ~10 % upon adding ATP to the imaging solution. Such multimerization via the globular domain has previously been found in the eukaryotic and archaeal MR complex and shown to promote endonucleolytic cleavage in DNA DSB repair (8, 9).

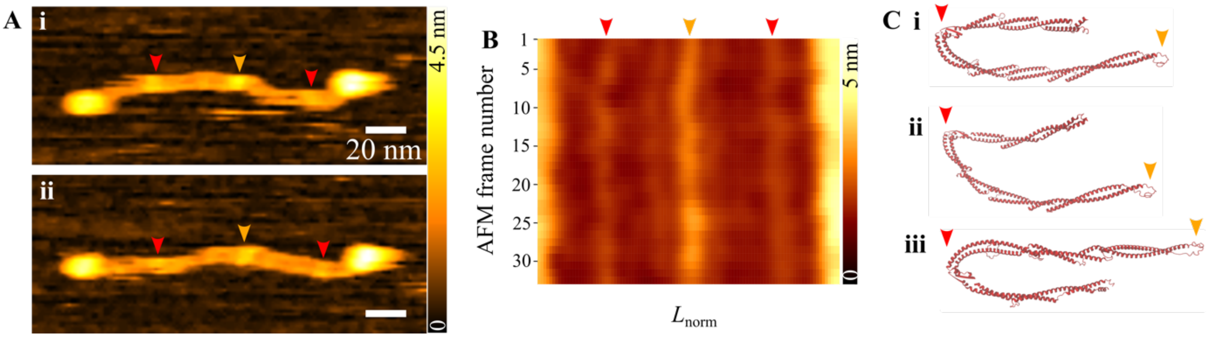


**Supplementary Figure S5. Insights into coiled coil structure. A. (i-ii)** Representative high-resolution HS-AFM topographs of the SbcCD dimeric coiled coil revealing three prominent high-contrast features along the contour length of the coiled coil, aside from those of the two globular domains at either end. The hook region is marked with an orange arrowhead, whereas the two other high-contrast features are marked with red arrowheads. The latter region most likely represents a local change in the packing of the monomeric coiled coil, resulting in its brighter appearance, and has shown bending from this region (see Supplementary Figure **S9**). **B.** A kymograph of the dimeric coiled coil shows variations in height along its contour length that remain spatially localized over time. The high-contrast feature in the middle corresponds to the Zn hook region characterized by a height of ~2.2 nm, located around 50 % of the distance from one of the globular domain ends. The other two features are located near the globular domains of each monomeric coiled coil segment and are likely due to differences in coiled coil packing, with a height of ~1.8 nm. These two features may be due to discontinuities of the heptad repeats leading to kinks or deformations along the coiled coils. **C. (i-iii)** A few AlphaFold2 predictions of the SbcC coiled coil (UniProt code: P13458) show folding (red arrowhead) within the coiled coil assembly, most likely due to discontinuities in the heptad repeats, consistent with the experimental observations.

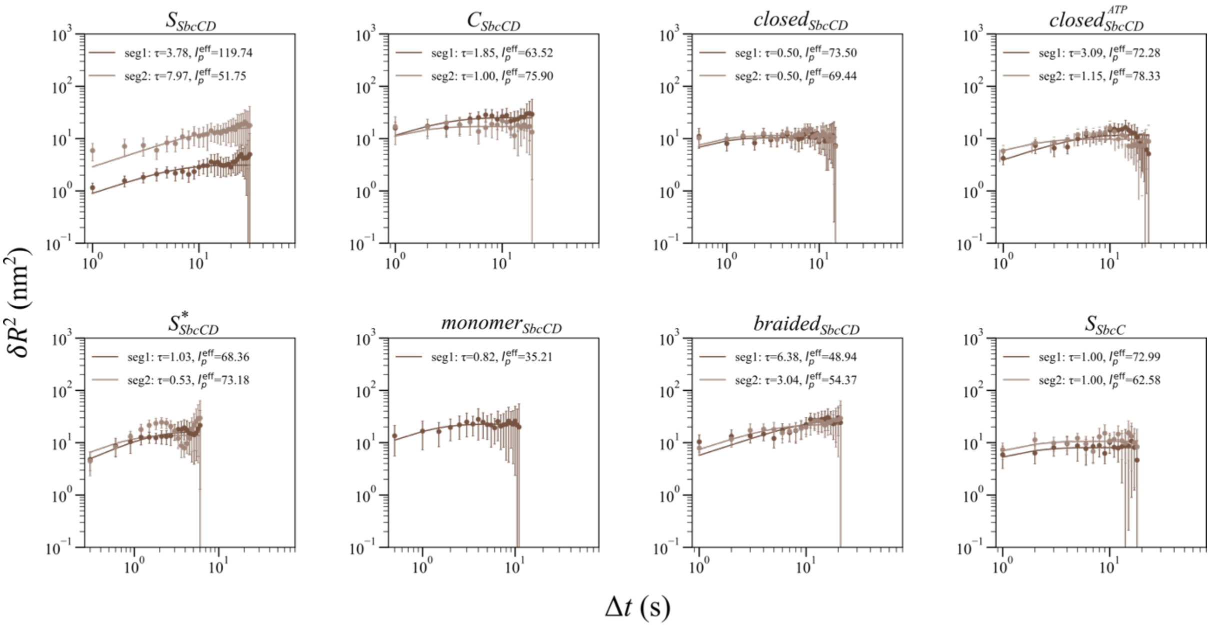


**Supplementary Figure S6. Determination of $l_p^{eff}$, $\zeta$, and $\tau$.** Representative plots of the MSD of the end-to-end distance ($\delta R^2$) from individual videos (mean ± SEM. The solid line shows the fit of **Eq. 1** from the main text, used to obtain the effective persistence length $l_p^{eff}$ (nm), internal friction coefficient $\zeta$ (pN·s/nm$^2$), and fundamental relaxation time $\tau_1$ (s) for various monomeric coiled coil segments (segments 1 and 2) across different conformations of the SbcCD complex and SbcC alone. The measured mean-squared deviation, $\delta R^2$, was fitted to the theoretical model described in **Methods** (Section 7), with $l_p^{eff}$ and $\tau_1$ as free parameters.

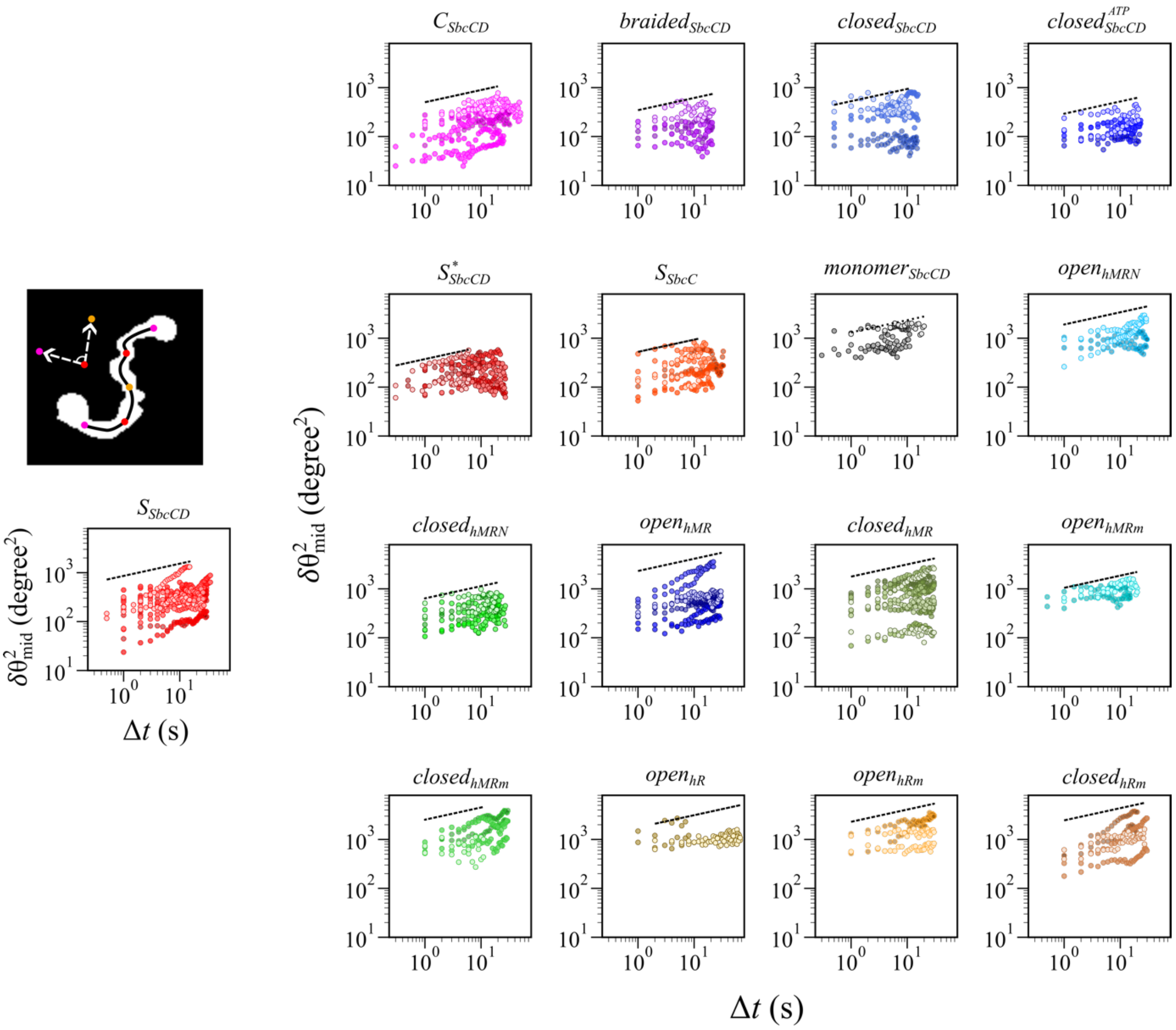


**Supplementary Figure S7. MSD of angle (θ). Left.** Binary image of $S_{SbcCD}$ with interpolated skeletal coordinates showing the angle determination to obtain the MSD of the angle for each monomeric coiled coil segment $\langle \delta\theta^2_{mid} \rangle$. **Right.** Plots showing $\langle \delta\theta^2_{mid} \rangle$ for each monomeric coiled coil segment across all the conformations of SbcCD, SbcC, hMRN, hMR, hMR$^{E1035\Delta}$, hR, and hR$^{E1035\Delta}$ as a function of elapsed time $\Delta t$ between HS-AFM frames. The dotted line represents a power law with a 1/4 exponent.

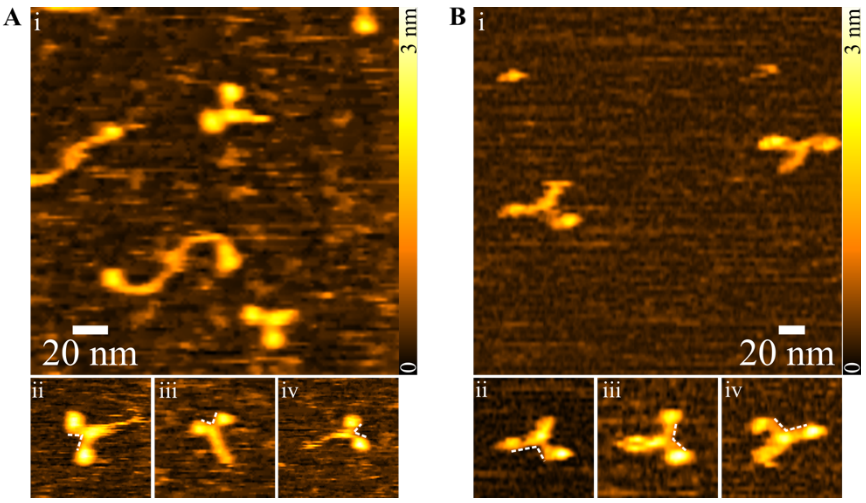


**Supplementary Figure S8. Braided conformations.** Large-area view of the two distinct $braided_{SbcCD}$ conformations of the SbcCD dimeric coiled coils based on the crossing region. In (**A-i**), the crossing point is in proximity to the globular domains, whereas in **(B-i)**, it is positioned approximately halfway along the contour length of each monomeric segment within the dimeric coiled coil, giving the structure a scissor-like appearance with the globular domains widely separated. The dynamic nature of both $braided_{SbcCD}$ is shown in **(A-ii–iv)** and (**B-ii-iv)**, with white dashed lines indicating the angles at the crossing regions. In **A**, the angle is < 90°, and > 90° for **B**. This variation in angle may have physiological relevance during the DNA-cutting state while enveloping the DNA. A similar rod-shaped structure has been reported from cryo-EM data of the *Ct*MRN dimeric coiled coil, where the globular domains were connected in the presence of ATP-γS (10).

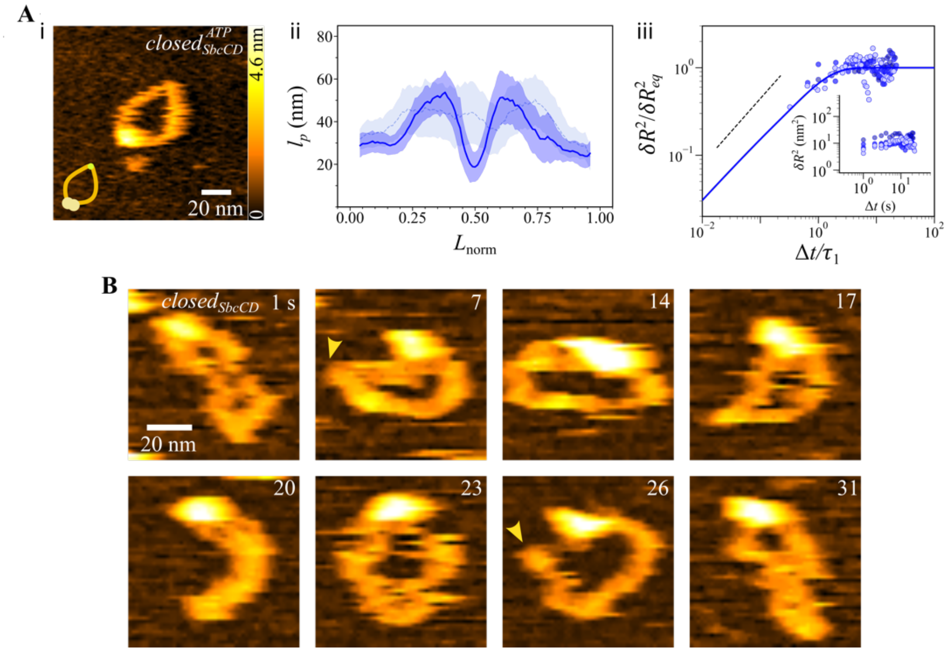


**Supplementary Figure S9. Closed conformation and the impact of ATP. (A-i)** A representative HS-AFM topograph of the $closed^{ATP}_{SbcCD}$ in the presence of ATP. Under this condition, we found more often $closed^{ATP}_{SbcCD}$ (~9 % as compared to ~6 % in the absence of ATP) where the two head domains were bound together and stabilized by ATP at their interface. **(A-ii)** Plot showing the local persistence length $l_p$ as a function of normalized contour length $L_{norm}$ for this conformation. For comparison, the $l_p$ of $closed_{SbcCD}$ in the absence of ATP is shown in its background. **(A-iii)** Rescaled MSD of end-to-end distance as a function of the rescaled time. The raw MSD data are shown in the inset. Overall, the ATP binding influenced the local flexibility and global dynamics of the coiled coil and, therefore, differed from the ATP-free state (see **Table 1**). **B.** Selected HS-AFM frames showing conformational changes (timestamp shown on the right side of each frame) in the $closed_{SbcCD}$ structure of the SbcCD dimeric coiled coils in the absence of ATP, captured at 1 fps. This conformation exhibited a high degree of structural flexibility and is known to be involved in genomic DNA scanning (11). We observed elbow formation along the contour length of the protein complex (indicated with yellow arrowheads). While elbow formation is well documented in the SMC family of proteins for DSB repair, it has not been extensively reported for the SbcCD complex (12). The elbow, which is most likely formed by changes in local packing within the coiled coil of the dimeric protein complex, as shown in Supplementary Figure **S2**, facilitates bringing the globular and hook regions of the complex into proximity. Such flexibility in the conformation may be essential for effectively encircling DNA during DNA sensing. Notably, unlike other conformations, the $closed_{SbcCD}$ and $closed_{SbcCD}^{ATP}$ conformation does not consistently preserve intrinsic curvature, giving rise to a range of closed-shaped intermediate conformations.

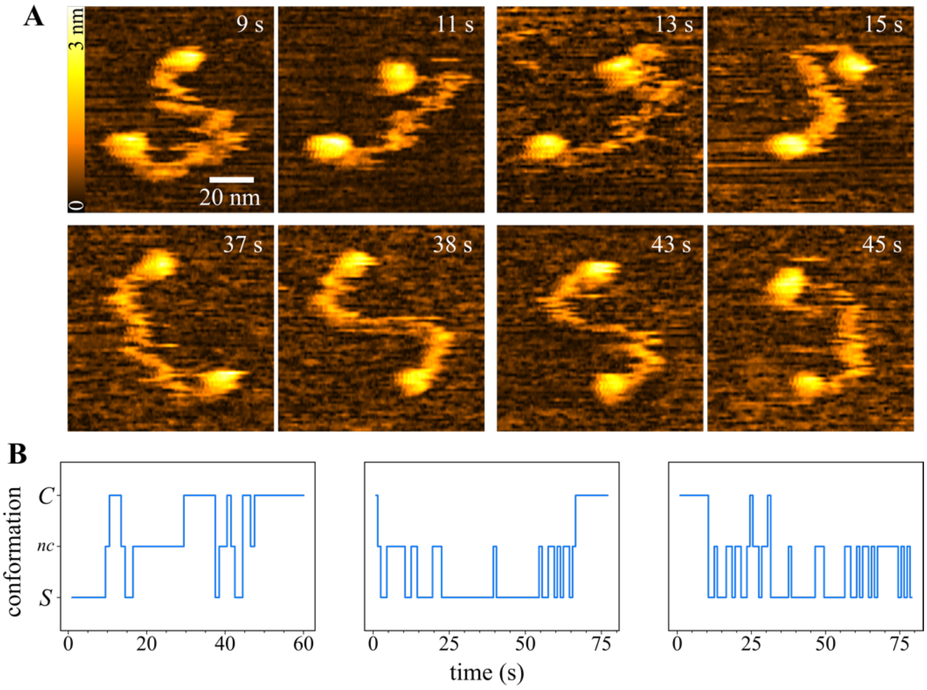


**Supplementary Figure S10. Conformational transition. A.** Representative HS-AFM frames showing the $S_{SbcCD}$-to-$C_{SbcCD}$/$C_{SbcCD}$-to-$S_{SbcCD}$ transition acquired at a scan rate of 1 fps, extracted from HS-AFM videos. Timestamps are shown in the top-right corner of each frame. **B.** Trajectories with such transitions from three independent HS-AFM videos. The conformations other than $S_{SbcCD}$ and $C_{SbcCD}$ in these cases are considered as non-classified ($nc$) conformations for the sake of clarity.

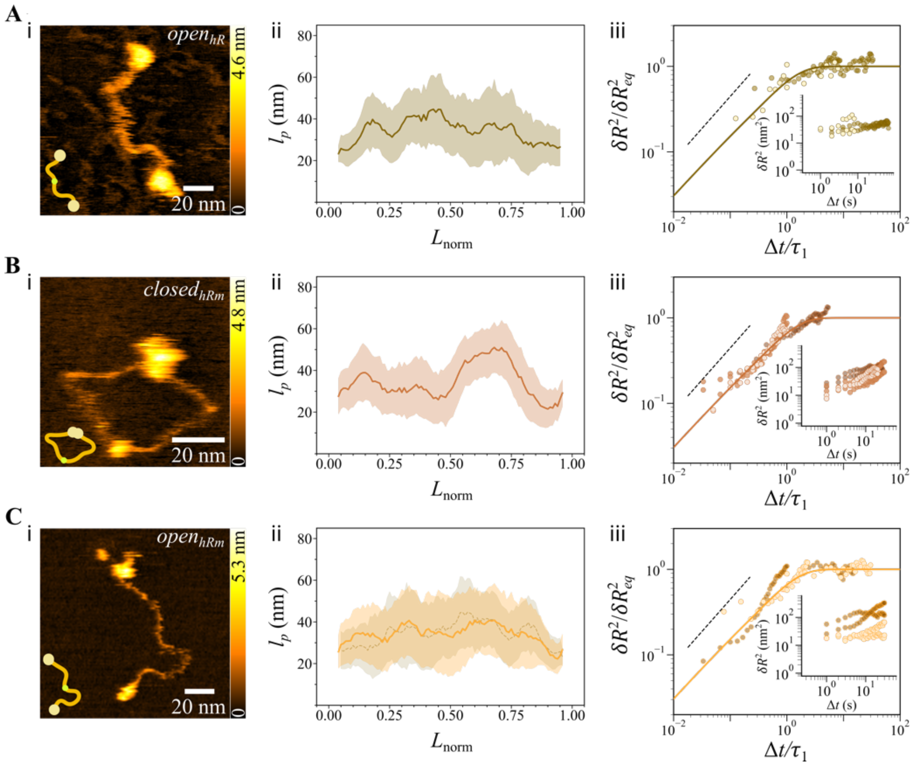


**Supplementary Figure S11. Human RAD50 conformations**. Representative HS-AFM topographs showing the different conformations of RAD50 alone for wild type and mutant: hR and hRm (or $hR^{E1035\Delta}$), respectively. (**A-i**) $open_{hR}$, (**B-i**) $closed_{hRm}$, (**C-i**) $open_{hRm}$ conformations. Plots showing the local persistence length $l_p$ as a function of normalized contour length $L_{norm}$ for the corresponding conformations: (**A-ii**) $open_{hR}$, (**B-ii**) $closed_{hRm}$, (**C-ii**) $open_{hRm}$. The $l_p$ of $open_{hR}$ is shown in the background of the $open_{hRm}$ for comparison. Rescaled MSD as a function of the rescaled time for the respective conformations: (**A-iii**) $open_{hR}$, (**B-iii**) $closed_{hRm}$, (**C-iii**) $open_{hRm}$. The raw data are shown in the insets of each rescaled plot.

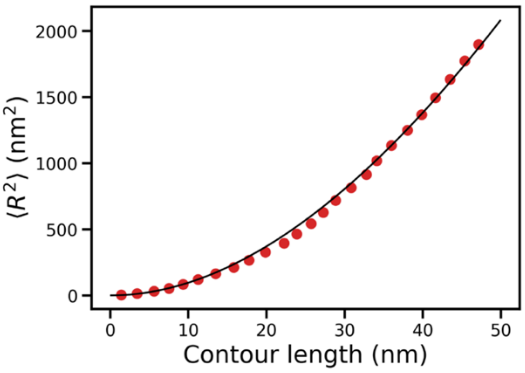


**Supplementary Figure S12. Two-dimensional end-to-end distance in MD simulations.** Average squared end-to-end distance as a function of contour length for the simulation of a coiled coil. Error bars associated with the points are smaller than the size of the symbols. The continuous line corresponds to the fit with the wormlike chain formula $\langle R^2 \rangle = 4 l_p L \left[1 - \frac{2l_p}{L}\left(1 - e^{-\frac{L}{2l_p}}\right)\right]$ with $l_p = 44$ nm.

#### Supplementary Tables

**Supplementary Table S1. HS-AFM videos used for analysis of flexibility ($l_p(L)$) and dynamics ($\langle \delta R^2 \rangle(\Delta t)$)**

| protein | conformation | no. of videos analysed | no of frames analysed | video with minimum no. of frames | video with maximum no. of frames |
|---|---|---|---|---|---|
| | $S_{SbcCD}$ | 8 | 429 | 28 | 70 |
| | $S^{*}_{SbcCD}$ | 5 | 238 | 40 | 55 |
| | $C_{SbcCD}$ | 6 | 293 | 39 | 65 |
| SbcCD | $braided_{SbcCD}$ | 3 | 129 | 42 | 45 |
| | $closed_{SbcCD}$ | 3 | 167 | 45 | 62 |
| | $closed^{ATP}_{SbcCD}$ | 3 | 135 | 43 | 47 |
| | $monomer_{SbcCD}$ | 4 | 174 | 40 | 45 |
| hMRN | $open_{hMRN}$ | 2 | 120 | 60 | 60 |
| | $closed_{hMRN}$ | 4 | 176 | 38 | 54 |
| hMR | $open_{hMR}$ | 4 | 222 | 45 | 60 |
| | $closed_{hMR}$ | 6 | 341 | 41 | 60 |
| $hMR^{E1035\Delta}$ or | $open_{hMRm}$ | 2 | 100 | 40 | 60 |
| hMRm | $closed_{hMRm}$ | 3 | 140 | 22 | 60 |
| hR | $open_{hR}$ | 2 | 82 | 16 | 66 |
| $hR^{E1035\Delta}$ or | $open_{hRm}$ | 2 | 120 | 60 | 60 |
| hRm | $closed_{hRm}$ | 3 | 154 | 40 | 60 |
| SbcC | $S_{SbcC}$ | 4 | 168 | 24 | 66 |

**Supplementary Table S2. HS-AFM video descriptions**

| Supporting Video (SV) | Description |
|---|---|
| SV1 | $S_{SbcCD}$ conformation |
| SV2 | $S_{SbcCD}$ high-resolution structure |
| SV3 | $S^{*}_{SbcCD}$ conformation |
| SV4 | $Z_{SbcCD}$ conformation |
| SV5 | $S_{SbcC}$ conformation |
| SV6 | $C_{SbcCD}$ conformation |
| SV7 | $S_{SbcCD}$-to-$C_{SbcCD}$ transition |

| | |
|---|---|
| SV8 | $braided_{SbcCD}$ conformation |
| SV9 | $closed_{SbcCD}$ conformation |
| SV10 | $closed_{SbcCD}^{ATP}$ conformation |
| SV11 | $monomer_{SbcCD}$ conformation |
| SV12 | $open_{hMRN}$ conformation |
| SV13 | $closed_{hMRN}$ conformation |
| SV14 | $open_{hMR}$ conformation |
| SV15 | $closed_{hMR}$ conformation |
| SV16 | $open_{hMRm}$ conformation |
| SV17 | $closed_{hMRm}$ conformation |
| SV18 | $open_{hR}$ conformation |
| SV19 | $open_{hRm}$ conformation |
| SV20 | $closed_{hRm}$ conformation |

**Supplementary Table S3. Mapping onto a coarse-grained polymer**

| coarse-grained bead | coiled coil | coil |
|---|---|---|
| 1 | 171-183, 897-904 | 171-183 |
| 2 | 184-196, 878-896 | 184-196 |
| 3 | 197-209, 864-877 | 197-209 |
| 4 | 210-222, 850-863 | 210-222 |
| 5 | 223-235, 840-849 | 223-235 |
| 6 | 236-248, 826-839 | 236-248 |
| 7 | 249-261, 800-825 | 249-261 |
| 8 | 262-274, 790-799 | 262-274 |
| 9 | 275-287, 763-789 | 275-287 |
| 10 | 288-300, 747-762 | 288-300 |
| 11 | 301-313, 734-746 | 301-313 |
| 12 | 314-326, 702-733 | 314-326 |
| 13 | 327-339, 688-701 | 327-339 |
| 14 | 340-352, 678-687 | 340-352 |
| 15 | 353-365, 668-677 | 353-365 |
| 16 | 366-378, 640-667 | 366-378 |
| 17 | 379-391, 626-639 | 379-391 |
| 18 | 392-404, 616-625 | 392-404 |
| 19 | 405-417, 610-615 | 405-417 |
| 20 | 418-430, 578-609 | 418-430 |
| 21 | 431-443, 564-577 | 431-443 |
| 22 | 444-456, 554-563 | 444-456 |
| 23 | 457-469, 543-553 | 457-469 |
| 24 | 470-482, 529-542 | 470-482 |
| 25 | 483-495, 523-528 | 483-495 |
| 26 | 496-522 | 496-503 |
